# Renewable energy caves: water replenishment holes in offshore monopiles create novel marine habitats

**DOI:** 10.64898/2026.06.03.729839

**Authors:** Leandra M. Kornau, Birte Leutelt, Christiaan J. van Sluis, Niek Bruinsma, Marjolijn D. Mascini, Renate A. Olie, Frank A.G. Jacobs, Sytske van den Akker, Eva de Haan, Eline van Onselen, Natalia Strigin, Reindert Nijland, Joop W.P. Coolen

**Author notes:** Corresponding author: LM Kornau. Author contributed equally (Birte Leutelt, Christiaan J. van Sluis).

## Abstract

The expansion of offshore renewable energy introduces artificial hard substrate, but the ecological effects may change depending on design features. For example, water replenishment holes, implemented for internal water refreshment and corrosion control, also allow colonisation of previously inaccessible monopile interiors, creating a novel semi-enclosed habitat. This study compared epifaunal communities on the interior and exterior walls of four water replenishment hole-equipped monopiles in the southern North Sea and modelled internal water quality to better understand factors shaping these communities. Vertical video transects were used to quantify percentage cover along depth gradients, while a coupled hydrodynamic-water quality model predicted vertical patterns in dissolved oxygen and particulate organic carbon over one year. Monopile interiors function as semi-enclosed, cave-like habitats with distinct environmental conditions, including darkness, restricted water flow, and vertical gradients in dissolved oxygen and particulate organic carbon. Compared to the external cover, interior communities showed reduced dominance of typical North Sea hard-substrate taxa and increased heterogeneity, with higher contributions of sponges, calcareous tube worms, and brittle stars, resembling communities reported from marine cave environments. Overall epifaunal cover was lower on the interior walls and broadly reflected vertical patterns in water quality. Organic matter accumulated on the interior seafloor, with indications of microbial mat formation. These findings suggest that the internal environmental conditions influence community development. Water replenishment hole design may therefore shape community composition inside monopiles, with implications for possible use as nature-inclusive design and environmental management.

**Graphical abstract:** 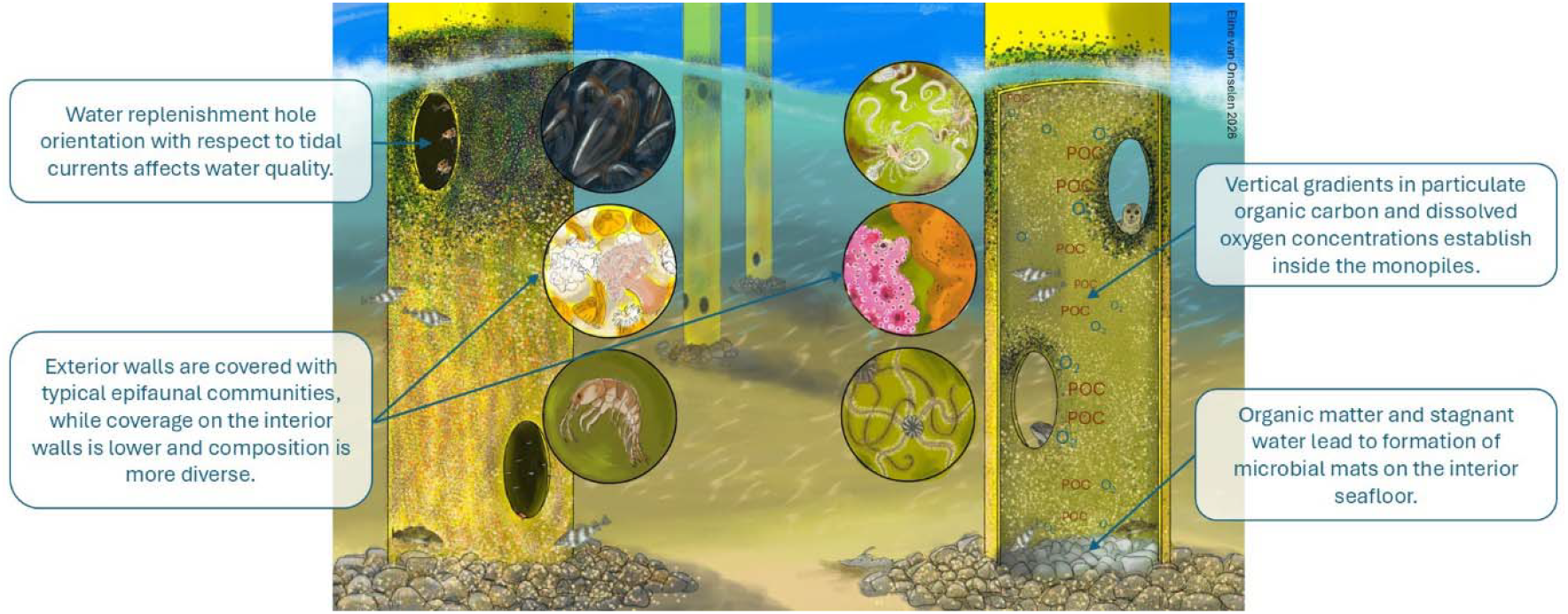

## 1. Introduction

The rapid expansion of offshore wind energy introduces large areas of artificial hard substrate (AHS) to marine environments (Degraer et al., 2020; Glarou et al., 2020; Van Sluis et al., 2025). These structures are quickly colonised by diverse epifaunal communities, increasing local biomass and habitat complexity (Coolen, Bittner, et al., 2020; Zupan et al., 2024). As a result, offshore wind farm monopiles have become key sites for studying colonisation on manl⍰made marine structures (Coolen et al., 2022; Degraer et al., 2020). Although distinct species zones establish along the water column (Krone et al., 2013), development of communities does not seem to reach a permanent climax situation over time (Zupan et al., 2023). The developing communities alter ecosystem functioning, including food web structure and biogeochemical cycling (De Borger et al., 2021; Mavraki, Degraer, Moens, et al., 2020). Understanding such patterns is increasingly important as offshore wind development reshapes coastal ecosystems, especially when introducing hard substrate in predominantly soft sediment habitats (Degraer et al., 2020).

New foundation features may further influence colonisation and thereby local biomass and diversity. An example of the last decade is the use of water replenishment holes (WRHs): openings in the submerged section of monopiles that allow passive exchange between internal and external seawater. WRHs and other larger openings have been integrated into North Sea monopiles in the Dutch North Sea since approximately 2019, with several wind farms (e.g., Borssele, Hollandse Kust Zuid and Hollandse Kust Noord) now employing this design feature. Their primary function is to mitigate the acidification of enclosed water and consequent risks of internal corrosion (Hilbert et al., 2011; Maher & Swain, 2018). Several studies on the technical aspects of their implementation have been published (Andersen, 2017; Carstensen, 2025; Jacobsen et al., 2025; Kesgin et al., 2024; Maher & Swain, 2018; Sarada et al., 2025; Tupkar & Sappe Narasimhamurthy, 2022). In addition to corrosion prevention, the openings also allow organisms to access and colonise the previously isolated internal hard substrate, effectively creating a novel habitat within the monopile (Maher, 2018). As WRHs become increasingly common in foundation design, more monopile interiors are converted into accessible habitat for epifauna and other demersal, mobile species.

In contrast to external monopile surfaces, which are exposed to light and strong tidal currents, monopile interiors represent semi-enclosed, dark environments. Within these environments, internal flow velocities can vary spatially and temporally depending on external hydrodynamic conditions and the size and number of WRHs (Jacobsen et al., 2025). Resulting reduced flushing may limit food resource and oxygen availability within the monopile. Despite increased application of WRHs, the ecological development within such man-made semi-enclosed habitats remains unknown and thus represents a critical knowledge gap in assessing the ecological impacts of offshore wind infrastructure. Studying hydrodynamics, water quality, and colonisation is highly relevant for understanding how WRHs influence biodiversity and ecosystem functioning. These insights can inform the development of offshore infrastructure design features that either enhance or mitigate ecological impacts, which is becoming increasingly relevant in offshore wind farm planning and tendering (Rijksoverheid, 2022; ter Hofstede et al., 2023) and international nature-positive ambitions (World Economic Forum, 2025).

This study characterises the epifaunal community developing on the interior walls of WRH-equipped monopiles and determines how internal environmental conditions and vertical gradients shape this community. We (1) used video footage to compare epifaunal percentage cover inside and outside monopiles along a vertical gradient two years after construction and to obtain an impression of the seafloor and (2) modelled vertical patterns in particulate organic carbon (POC) and dissolved oxygen (DO) concentrations inside a monopile over one year.

## 2. Materials & Methods

### 2.1. Site description

This study was conducted in the offshore wind farm Hollandse Kust Zuid (HKZ), located at a minimum distance of 18 kilometres from the Dutch coast (Fig. 1). The dominant sediment type in this region is sand, with large and medium sand dunes moving through the area (Fugro Survey B.V., 2016). The water column is influenced by freshwater input from the Rhine, which can produce salinity stratification, while thermal stratification is negligible (Van Duren et al., 2021). The four monopiles investigated in this study (D1, E5, F1 and F4) are all located in the northwestern part of the wind farm. They were installed between July and August 2021 and are situated at depths from -23.26 m to -25.24 m (Table S1).

**Fig. 1:**
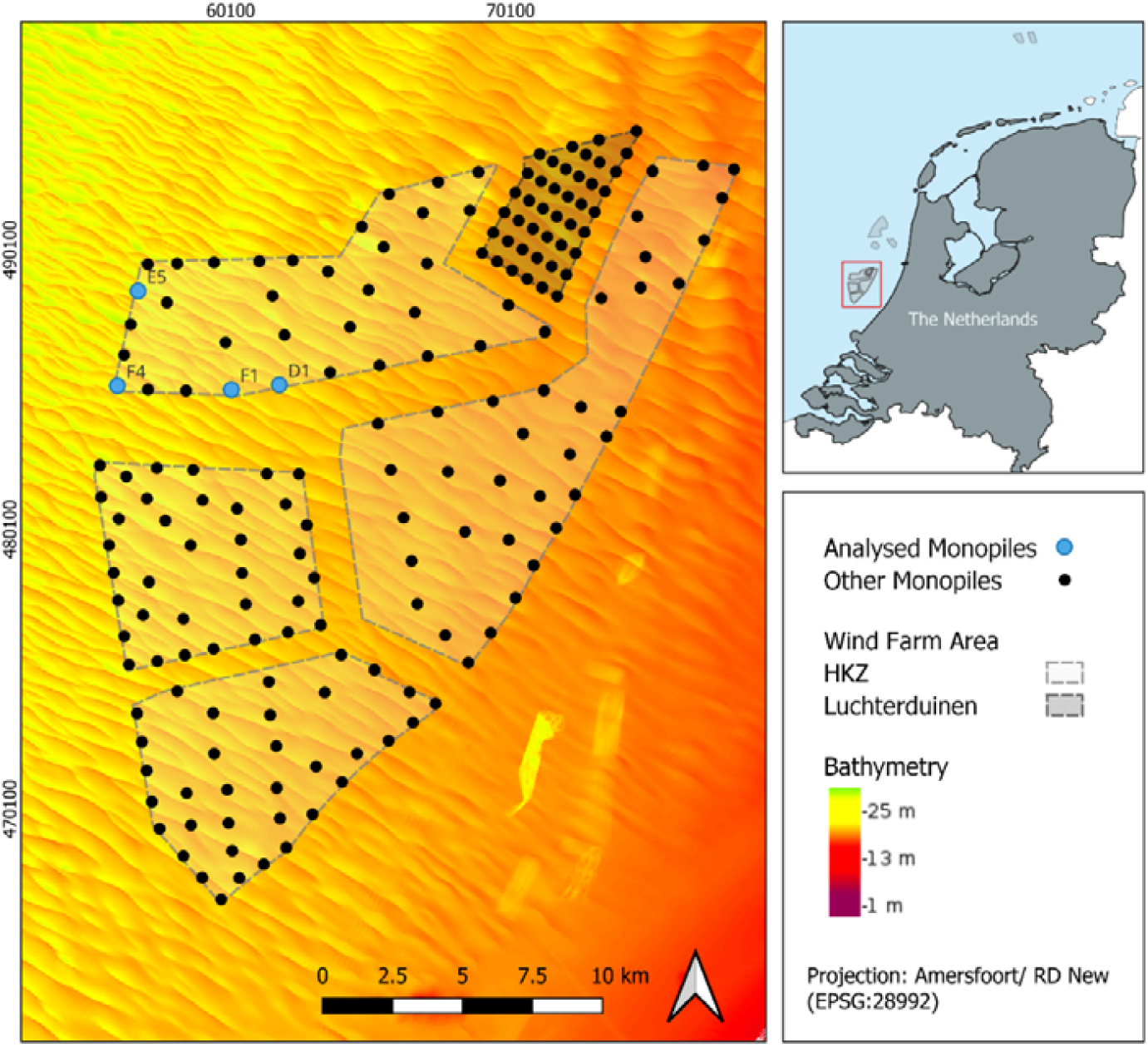
Map of the offshore wind farm Hollandse Kust Zuid relative to the Dutch coast and highlighting monopiles D1, E5, F1, and F4 (Global Administrative Areas, 2025; Rijkswaterstaat, 2024; Rijkswaterstaat & Deltares, 2024). Created in QGIS 3.4.11.

### 2.2. Water replenishment holes

Each monopile is equipped with four elliptical WRHs of 0.96 m height and 0.32 m width. The positions of the holes were adapted for each monopile, so that two are located at an average height of 4.35 m above the seabed and two 1 m below the water surface (lowest astronomical tide, LAT). The upper WRHs have a fixed orientation of 191°N and 255°N. The lower WRHs were oriented at 191°N and 258°N for most investigated monopiles, except for monopile F1, which had the lower WRHs oriented at 191°N and 25°N.

### 2.3. Biotic and abiotic field measurements

Several environmental parameters were measured on the outside and inside of monopile F1 on 16 October 2024 (Table S2). Two Eureka Manta 35+ multiprobes equipped with sensors for DO, chlorophyll-a, turbidity, temperature, pH, conductivity, and depth were used with a 0.5 Hz sampling interval. At three different depths (upper and lower WRH, and middle of the monopile), parameters were monitored for 10 minutes. This monitoring was executed twice both inside and outside. Abiotic data is provided in Supplementary Data 2.

Vertical transect video recordings of the exterior monopile surfaces were collected between 30 August and 9 September 2024 (Table S2) using a remotely operated vehicle (ROV, Saab Seaeye Falcon®), around neap tide conditions to ensure stable operation and consistent image quality. A forward⍰facing camera (GoPro) was mounted on the ROV at a slight downward angle, recording at 59,94 fps and with *“*wide view*”*. Two parallel laser pointers spaced 27.5 cm apart were used to estimate the surface area visible in the recordings. Interior recordings were collected using a mini⍰ROV (VideoRay Pro 4) equipped with a camera (GoPro) mounted on top. Laser scaling devices were not available on the mini⍰ROV, preventing estimation of the visible surface area in individual still frames. Nine exterior and sixteen interior transects were recorded (Table S3). In addition to vertical transects, seafloor footage was also collected using the same ROV systems. These recordings were used to qualitatively assess seabed composition and organic matter accumulation on the interior and exterior of the monopile. Recordings and selected frames are available online under https://doi.org/10.5281/zenodo.20508299

### 2.4. Video analysis

#### 2.4.1. Frame selection

For each vertical transect, approximately one frame per depth meter was manually selected from the recorded footage when image quality was sufficient for taxonomic identification. In total, 317 interior frames and 157 exterior frames were analysed (Table S3). Due to the different natural light conditions, the overall quality of the interior frames was generally lower. Therefore, frames closer to the wall needed to be chosen to provide the best image quality, which resulted in smaller surface areas analysed for the interior frames.

#### 2.4.2. Taxonomic classification and percentage cover

For each frame, a rectangular area of interest (AOI) was defined only including image regions with sufficient quality for taxonomic identification, excluding blurred regions and water. AOI size was estimated based on the vertical and horizontal camera angle and object-scale references derived from the organism size within the image. A point grid (200 points/m^2^) was overlaid using TransectMeasure (SeaGIS-V4.23) and AOI size included as covariate (section 2.5) to account for the differences in AOI between frames (Fig. S1). Each point was classified as the lowest identifiable taxonomic group, *“*not colonised*”* (bare substrate or biofilm), or *“*blurry*”* when identification was not possible due to hyperlocal low image quality. Blurry points (10.5 % of the total) were excluded from further analysis.

The percentage cover per taxon was calculated for each frame based on the remaining points. The average percentage cover per taxonomic group per depth was calculated across transects inside and outside for each monopile. All statistical analyses were carried out using R (R Core Team, 2025) in RStudio (version 2024.04.2 Build 764; Posit Team, 2025). Visualisations were done using the ggplot2 package (Wickham, 2016).

#### 2.4.3. Biomass estimation

To derive biomass estimates along the depth gradient for use in the water-quality model, a semi-quantitative approach was applied to the frames of monopile F1. Frames were first visually ranked within each transect from lowest to highest biomass based. For the exterior frames, the highest and lowest biomass frames were compared with reference footage from the HKZ-E4 marine growth sampling tool campaign (November 2023; Coolen et al., 2025) and from the gas platform L10-AD (October 2015; Coolen, Van der Weide, et al., 2020). When similar species composition and biomass were observed, corresponding ash-free dry weight (AFDW, g m^-2^) values of the collected scrape samples were assigned. For platform L10-AD, wet weight values were first converted to AFDW using species-specific conversion factors derived from published and internal laboratory datasets (Dannheim, Kloss, et al., 2025; Wageningen Marine Research Den Helder, unpublished data).

On the interior of the monopile, no suitable reference footage was available due to differences in community structure with any available data. Therefore, individual organisms were counted in lowest and highest biomass frames and converted to AFDW using species-specific mean individual weights, scaled to frame area. Linear interpolation between ranked frames was then used to assign AFDW values to all remaining frames along each transect.

### 2.5. Statistical analyses

To generalise vertical patterns in species cover percentages observed inside and outside across monopiles, generalised linear latent variable models (GLLVMs) were created using the gllvm package (Niku et al., 2019; Van der Veen, 2025). GLLVMs provide statistical testing of relations between multiple environmental variables and multiple species simultaneously. Model preparation followed Zuur & Ieno (2025) and van der Veen (2025). Only species with a minimum of 15 presence observations in the full dataset were included in the model.

Model formulation took the form of a polynomial regression with quadratic and cubic terms for depth, in interaction with a factor variable inside versus outside, to account for differences in depth patterns. The estimated AOI of each frame was included to account for differences in area analysed (Fig. S1). The inclusion of a parameter on the modelled oxygen concentration was considered, but both mean and minimum oxygen concentration were highly collinear with depth (Fig. S3). Therefore, only depth was included, as depth-related patterns on the outside of monopiles have been reported multiple times (Coolen et al., 2022; Dannheim, Beermann, et al., 2025; Krone et al., 2013) and as depth is a proxy for water quality for the inside.

Latent variables were included to account for relations with unknown environmental variables not incorporated in the model. To test whether latent variables were essential, models with 0, 1 and 2 latent variables (*jitter*.*var* set to 0.1, with 25 initial runs per model) were created and compared using the Akaike Information Criterion (AIC) (Akaike, 1973). The model with 2 LV had the lowest AIC and was used to present further results. Video transect ID nested in turbine ID was included as a random effect to compensate for dependency between observations caused by repeated sampling of the same monopile. Large numbers of zero observations were present in the data of most species. Therefore, the data were assumed to be zero-inflated, and a beta hurdle distribution (betaH) was applied (Korhonen et al., 2024), using the extended variational approximations (EVA) approach (Korhonen, 2025; Korhonen et al., 2023).

Model validation was performed by plotting residuals against fitted values, variables included in the model, and fitted vs observed values, inspecting the plots visually to confirm assumptions of homogeneity of variance and normality were met. Generalised species pattern plots to visualise differences between the interior and exterior of monopiles along the depth gradient were created using data from the predict.gllvm function. For prediction purposes, the estimated AOI was set to the average in the complete dataset. Model output is available in Supplementary Data 4.

### 2.6. Numerical modelling framework

Abiotic conditions inside offshore wind monopile foundations equipped with WRHs were quantified using a coupled numerical modelling framework (Supplementary Data 1 section 1.2). The framework combines a hydrodynamic model that translates external metocean forcing into exchange fluxes through WRHs with a onel⍰dimensional vertical (1DV) water quality model that simulates internal distributions of DO and POC. The water column inside the monopile is divided into 61 layers, assuming horizontal homogeneity. Exchange with the surrounding water is represented as flux through the WRHs driven by external hydrodynamic forcing. The model is forced using wave, current, and water level data with hourly resolution, and subsequently estimates the local hydrodynamics and water quality at a temporal resolution of 1 s. Vertical advection and diffusion were included as internal transport processes, while biogeochemical dynamics are represented by POC sedimentation and biotic consumption of DO and POC.

Ambient concentrations of DO and POC were prescribed as boundary conditions for inflowing water. Oxygen concentrations were assumed constant and representative of offshore North Sea conditions (8.0 mg/L). Ambient POC concentrations (0.42 mg/L) were estimated from measured chlorophyll concentrations outside the monopiles using fixed conversion ratios. Initial internal concentrations were set equal to ambient values. Model performance was evaluated against in situ measurements (Fig. S5). The model reproduces the observed order of magnitude and the vertical structure of DO and POC, including elevated concentrations near WRH elevations and reduced concentrations in poorly flushed zones.

## 3. Results

### 3.1. Community composition and cover patterns

In total, 28 taxa, comprising 9 phyla, were identified. Differences in community composition were observed between interior and exterior monopile surfaces as well as along the vertical gradient.

While exterior surfaces were almost completely colonised (97.4 ± 4.3 %, indicated as mean cover ± standard deviation in this section), the interior walls exhibited more bare area (84.4 ± 15.1 %). The total cover on the interior of the monopiles was generally higher near the water surface and seafloor and lower in middle range depths (around 8-16 m; Fig. 2).

**Fig. 2:**
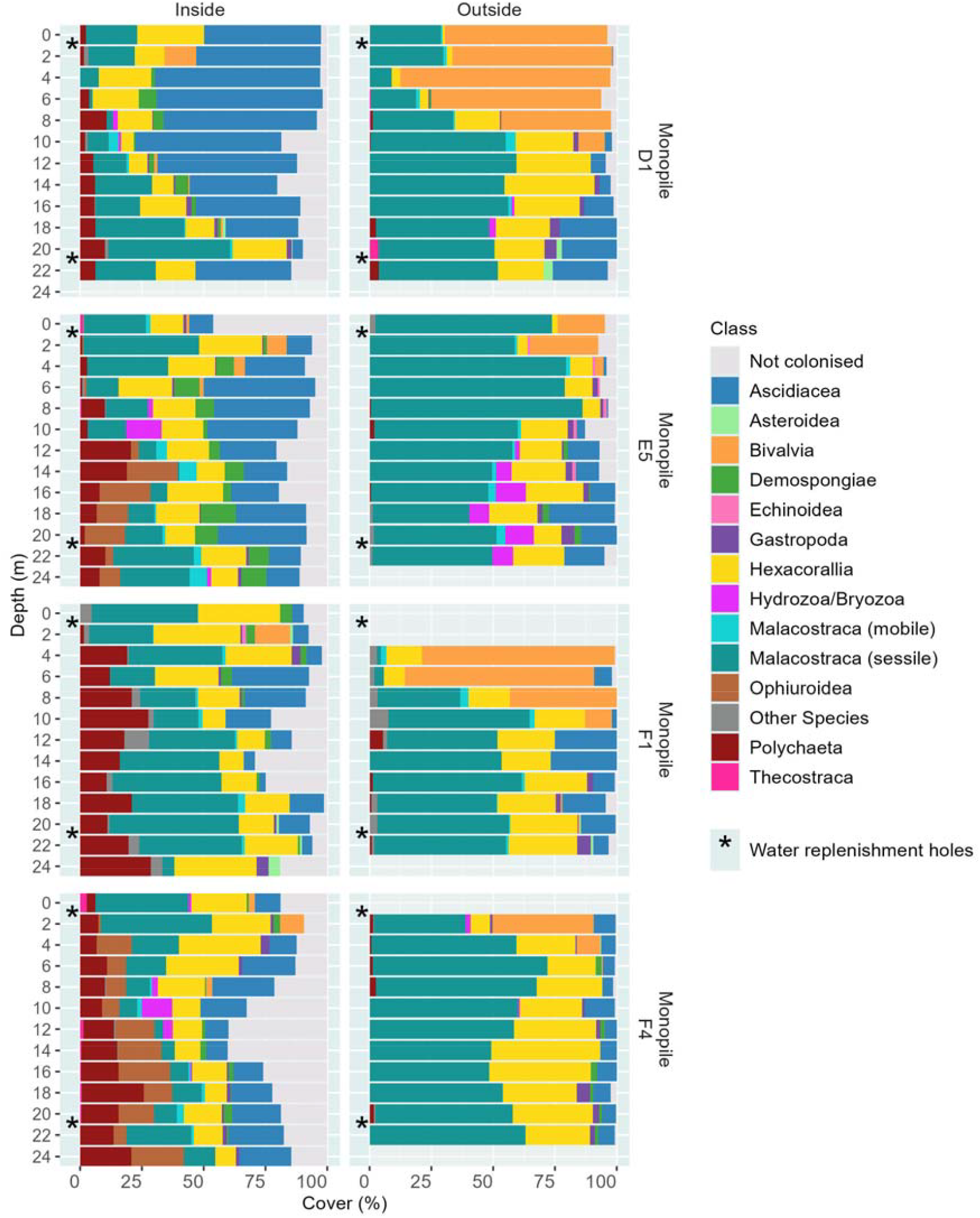
Mean percentage cover of the exterior and interior of monopiles D1, E5, F1, and F4 per taxonomic category for different depths. * indicates the vertical position of the water replenishment holes. Category *‘*Other species*’* includes unidentified species, *Alcyonium digitatum*, and *Ulva* spp. (the latter with two observations). Underlying data is available in Supplementary Data 3.

On the exterior of the monopiles, mats assumed to be built by Amphipoda (sessile Malacostraca), anemones (Hexacorallia), Bivalvia, and colonial Ascidiacea were dominant, collectively accounting for an average of 78.1 ± 12.9 % of the total cover. Bivalvia were particularly dominant between 0 and 8 m on monopiles F1 and D1, but this zone was restricted to 0-4 m for monopiles F4 and E5. Most Bivalvia were blue mussels (*Mytilus edulis*) however on monopile F4, the non-indigenous Pacific oysters (*Magallana gigas*) dominated. Below the Bivalvia-dominated zones, Amphipoda became more prevalent, combined with varying covers of different Hexacorallia species and colonial Ascidiacea (mainly *Diplosoma listerianum* species complex).

On the interior walls, Bivalvia were largely absent. *M. edulis* occurred only in the immediate vicinity of the upper WRHs and *M. gigas* were observed sporadically. Similar to the communities in the deeper parts of the exterior of the monopiles, the communities on the interior consisted of Amphipoda mats, colonial Ascidiacea, and different Hexacorallia species, but these dominant exterior species had lower cover on the interior (54.6 ± 23.0 %). Layer thickness (not measured) of these species also appeared to be much lower on the interior walls. In contrast to the exterior, additional taxonomic groups reached higher cover percentages on the interior, including Porifera (Demospongiae), brittle stars (Ophiuroidea), and Serpulidae (Polychaetae). Specifically, Serpulidae consistently covered a large part of the walls, with the highest cover (up to 39 %) at intermediate depths. Ophiuroidea were exclusively observed on the interior of the monopiles. The non-indigenous slipper limpet (*Crepidula fornicata*, Gastropoda) had a low cover on the interior (0.7 ± 1.9 %) compared to the exterior (1.5 ± 2.8 %).

Differences between monopiles were also evident on the interior walls. On the interior of D1 and F1, no Ophiuroidea were observed. Colonial Ascidiacea covered less area on the interior of F1 and F4. The overall area covered by epifauna on the interior of this monopile was higher than for the other monopiles. Porifera were particularly prevalent in monopile E5. These observed differences were further examined using species-environment models.

### 3.2. Species-environment relationships

The predicted taxa-specific trends between interior and exterior surfaces as well as along the depth gradient were assessed using the GLLVM (Fig. 3). Significant differences (all reported when P<0.05) in observed cover percentage between the interior and exterior of the monopiles were present in four taxa (Supplementary Data 4). Colonial Ascidiacea and Serpulidae percentages were significantly higher on the interior than on the exterior, while *M. edulis* and Amphipoda percentages were significantly lower on the interior than on the exterior.

**Fig. 3:**
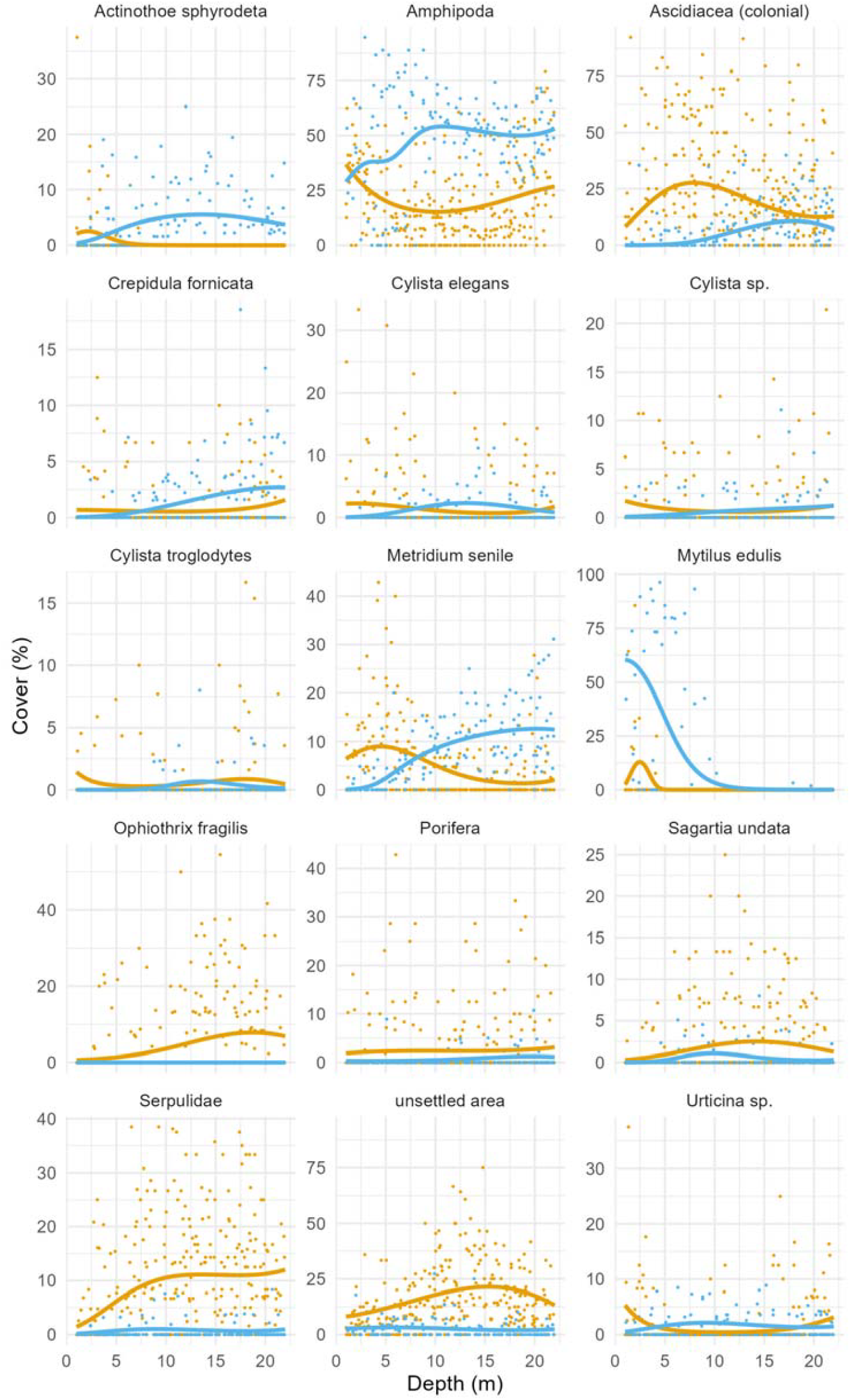
Generalised linear latent variable model results per taxon for interior (orange lines) and exterior (blue lines) of the monopiles, along the depth gradient, with observed data (dots) from all four monopiles).

In addition to the percentages, taxa presence likelihood (predicted within the hurdle part of the model) of *Sagartia undata* was significantly lower on the exterior than on the interior, while likelihood of *Metridium senile, C. fornicata, Cylista elegans* and *Urticina* sp. was significantly lower on the interior than on the exterior.

Depth patterns could be visualised for a number of taxa (Fig. 3). Significant differences in percentage cover were observed in 4 out of the 19 taxa for the linear depth-cover relation, 5 for the quadratic term and 6 for the cubic term. *M. edulis* was most abundant in shallow parts of the exterior walls, a pattern significantly different from the interior. In deeper sections, the species showed low percentages on both the interior and exterior. Amphipoda had similar cover percentages on the interior and exterior at shallow depth, but at depths from mid-water to near-seabed, percentage cover was significantly lower on the interior than on the exterior. Colonial Ascidiacea showed a significant depth-related pattern, with the highest cover percentages in shallow to mid-water sections of the interior of the monopile and with lower percentage cover on the exterior. Serpulidae, while near-absent on the exterior, showed increasing percentages with depth on the interior. *M. senile* showed opposing patterns on the interior vs the exterior, with high densities in shallow water on the interior, reducing with increasing depth.

### 3.3. Organic matter accumulation

In addition to the vertical patterns, clear differences in seafloor structure were observed between the interior and exterior of the monopiles. The seafloor on the interior was largely covered by soft material, present on top of the scour protection layer (Fig. 4). The distribution of rocks and soft substrate was patchy. Grey layers on the seafloor, interpreted as microbial mats forming under hypoxic conditions, were present on the interior of all four monopiles. In contrast, exterior recordings showed that the original rock substrate was not covered by sediment or loose organic matter, with only spaces in between rocks filled with sediment and no microbial mats.

**Fig. 4:**
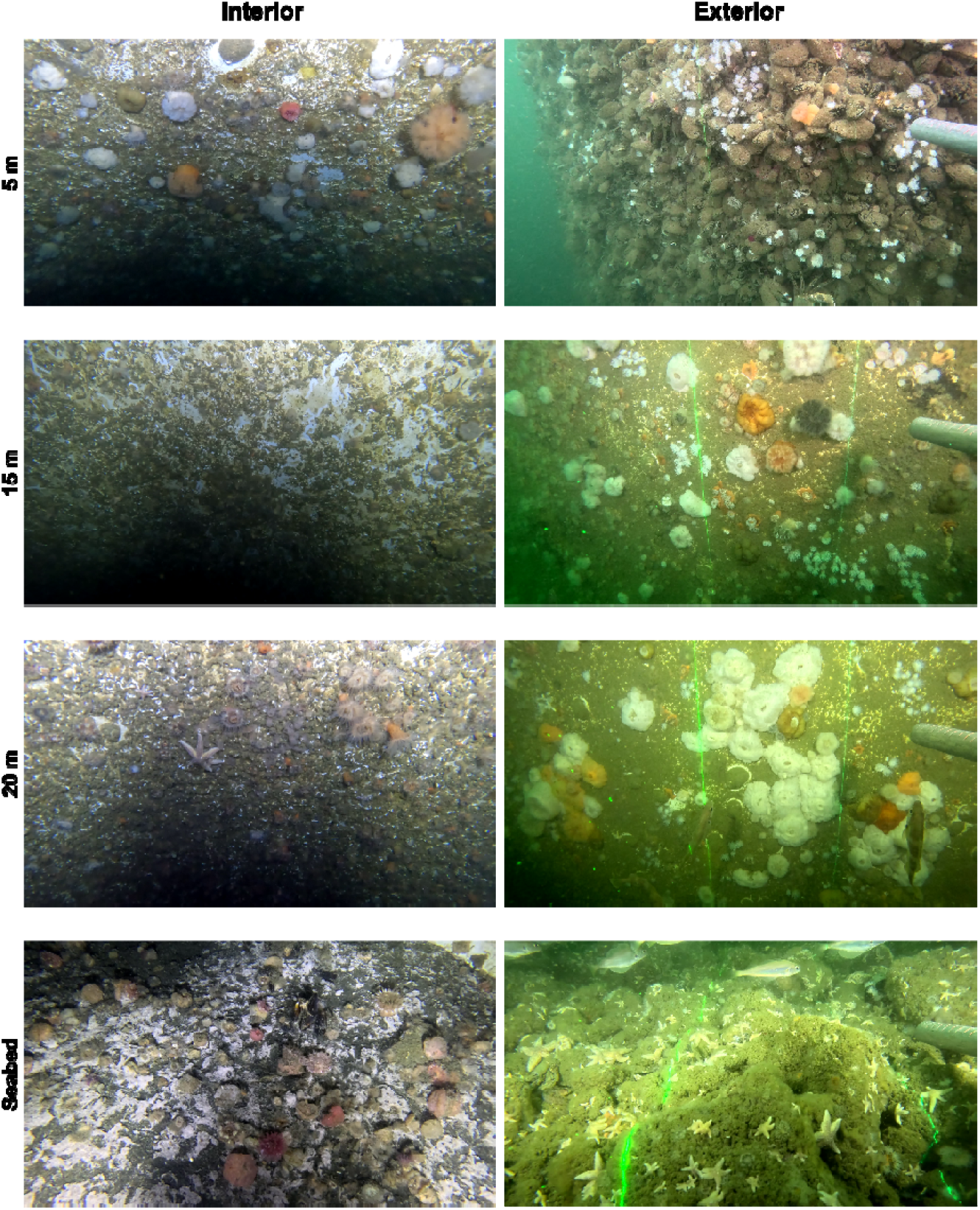
Communities at different depths on the vertical walls and seabed on the interior and exterior of the monopiles. The visible amount of organic matter on the interior of the monopiles is higher than on the exterior. The grey layers likely are microbial mats forming under hypoxic conditions.

### 3.4. Modelled environmental conditions

The numerical model simulated the spatial and temporal variability of DO and POC on the interior of the monopiles (Fig. 5). Both variables exhibited a pronounced vertical structure. Along the vertical profile, concentrations were elevated in the vicinity of the WRHs and reduced in zones above, below and between the WRHs. This vertical gradient reflects differences in replenishment efficiency, as exchange with ambient seawater is strongest near the openings and weaker in more isolated parts of the water column. The clear temporal variability follows temporal changes in external hydrodynamic forcing. Periods with increased wave activity and stronger currents correspond to enhanced water exchange through the WRHs and lead to higher internal DO and POC levels and reduced vertical gradients. Conversely, during calmer periods, replenishment was weaker and vertical gradients became more pronounced, particularly in the lower part of the monopile. These patterns indicate that internal environmental conditions are strongly controlled by external forcing, with waves playing a key role in short-term variability and seasonal changes in exchange intensity.

**Fig. 5:**
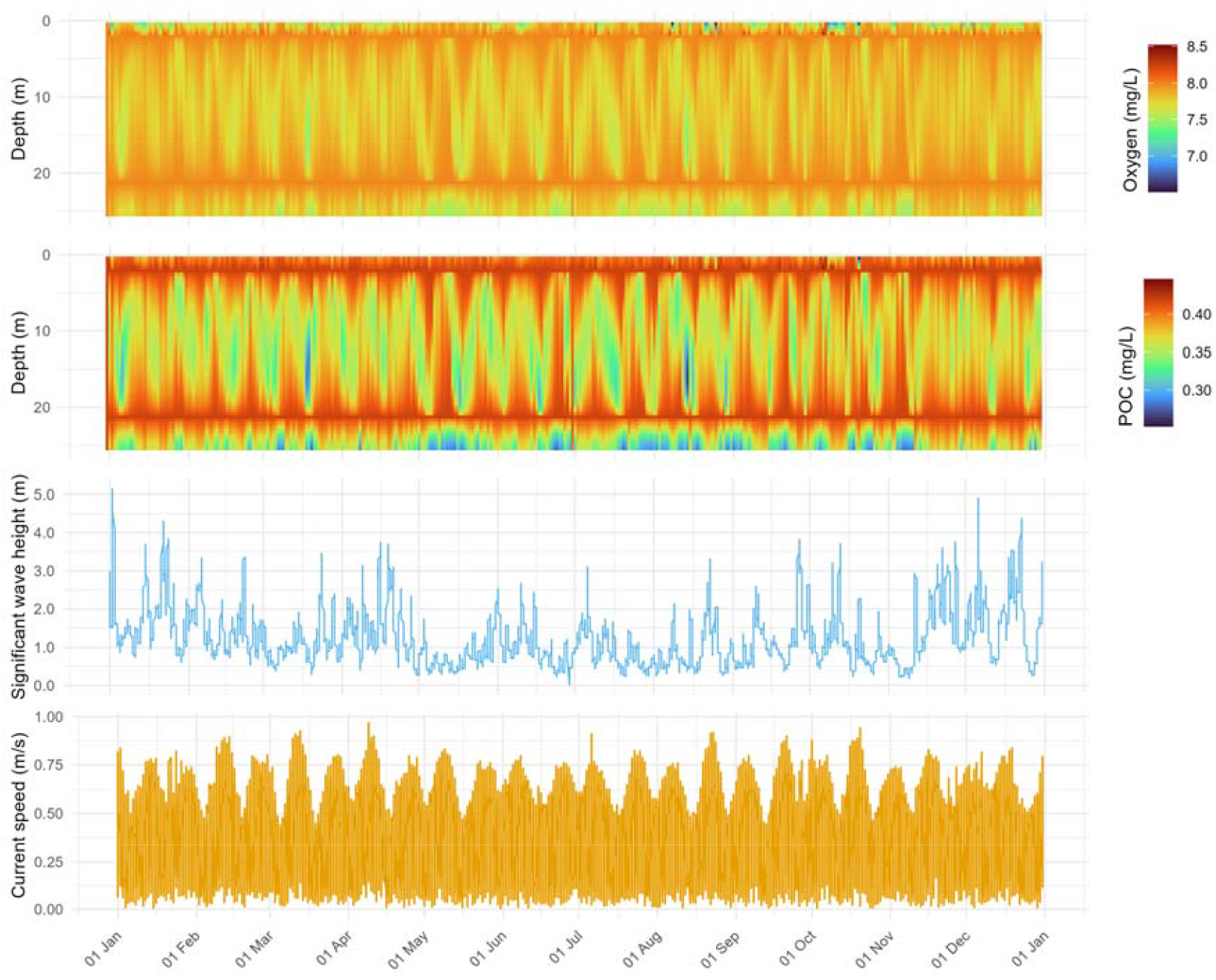
Hourly modelled DO (top), POC (middle), and ambient hydrodynamic conditions: spectral wave height and absolute current velocity (bottom) for the F1 monopile for 2024.

Modelled mean concentrations of DO and POC were similar for monopiles D1, E5, and F4, which share the same WRH configuration (Fig. 6). Differences between these monopiles are minor and primarily related to variations in water depth, which slightly affect pressure forcing and internal volume. Monopile F1 showed higher mean concentrations and a reduced vertical gradient. Fig. 6 C and D illustrate instantaneous modelled conditions on 20 March 2024 corresponding to the moment of maximum inter-monopile differences in DO and POC. In monopile F1, the lowest DO and POC concentrations occurred near the lower WRHs rather than between the upper and lower WRHs. Taken together, the model results demonstrate how internal oxygen and POC conditions are governed by the interaction between WRH configuration, monopile geometry, and external hydrodynamic forcing.

**Fig. 6:**
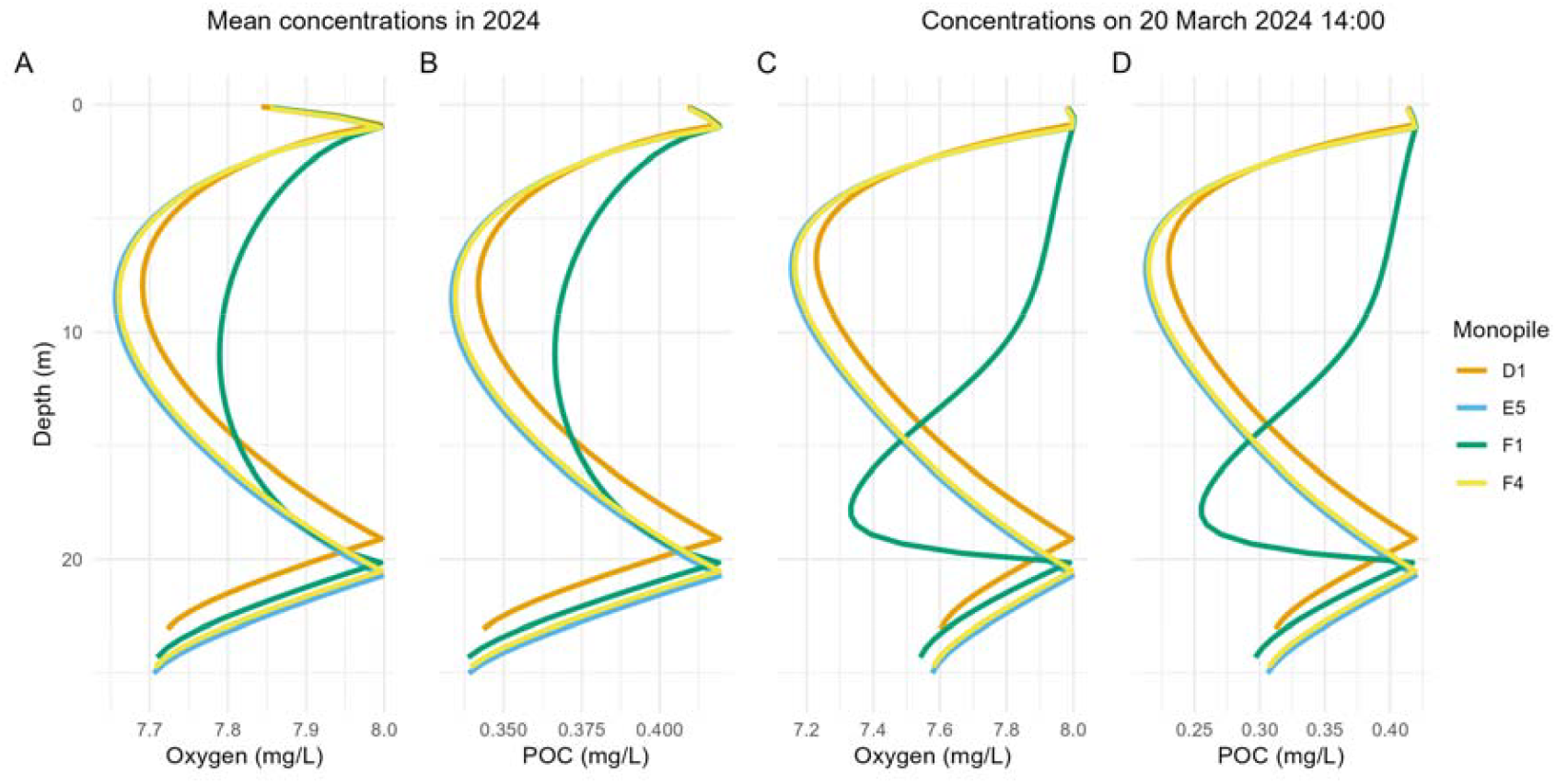
Mean concentrations for 2024 of A: Dissolved oxygen (DO); B: Particulate organic carbon (POC). Concentrations on 20 March 2024 for C: DO and D: POC for the four different monopiles.

## 4. Discussion

This study provides the first investigation of the water quality and epifaunal communities developing within the interior of offshore wind monopiles equipped with WRHs. The results demonstrate that within these monopiles a specific semi-enclosed habitat develops with epifaunal communities distinct from the exterior walls. These communities differ in composition and total cover percentage over the water depth gradient.

Modelled DO and POC concentrations near the WRHs resembled the surrounding water masses and are reduced around mid-depth, between the WRHs, and below the lower and above the top WRHs. Patterns in the epifaunal community composition and total cover reflect these gradients, with the lowest cover in the mid-depth section on the interior walls. The interior communities also generally exhibited lower overall cover, reduced dominance of typical exterior taxa such as Amphipoda, Hexacorallia, and Bivalvia, and increased contributions from taxa such as Porifera, Serpulidae, and Ophiuroidea. A comparison of the four studied monopiles indicated differences in community composition among monopiles and that the orientation of the WRHs mattered for the interior water quality. Together, these findings suggest that internal environmental gradients may influence epifaunal community composition and that the implementation of WRHs creates a novel ecological habitat within offshore wind farms.

### 4.1. Exterior and interior communities

Artificial hard substrate in the southern North Sea typically shows depth-driven vertical zonation patterns in the epifaunal communities: a mussel-dominated layer in the upper 5 m, an Amphipoda-dominated mid-layer (*Jassa herdmani*, 5-15 m), and a layer of plumose anemone (*Metridium senile*) extending towards the seabed (Coolen et al., 2022; Krone et al., 2013). The communities on the exterior of the investigated monopiles broadly corresponded to these general patterns, with a Bivalvia-dominated shallow zone and Amphipoda and Hexacorallia present in deeper zones. However, we did not observe a clear distinction between Amphipoda- and Hexacorallia-dominated zones.

The interior walls exhibited different patterns. Specifically, the absence of Bivalvia except near WRHs, the increased presence of Porifera and Serpulidae, and new appearance of Ophiuroidea, which were not observed on the exterior walls, stand out. Increased presence of Serpulidae was also observed inside enclosed space in a shipwreck (Walker et al., 2007). Several mechanisms may explain the observed patterns, including environmental stress, species*’* interactions, and larval settlement. The differences are consistent with the variation in environmental conditions between the interior and exterior as well as along the vertical axis. Reduced DO and POC, limited water flow, and constant darkness create environmental stress inside the monopile that may constrain taxa commonly found on AHS. Although DO is only marginally reduced and still within normal limits, the reduction in flow speed and POC concentration are likely explainers for the observed community shift (Riisgård & Larsen, 2015; Wildish & Kristmanson, 1997). Exterior wall communities are likely shaped through an inhibitory response (Connell & Slatyer, 1977; Coolen et al., 2022), meaning that a diverse colonisation is prevented by early colonisers with strong competitive traits. *M. senile* and Jassa spp. are extremely successful colonisers as they prevent settlement of other species through e.g., occupying space, overgrowing or smothering recruits, and feeding on other species*’* larvae (Beermann, 2013; Nelson & Craig, 2011). In our observations, *M. senile* cover on the interior of the monopiles was highest near the water surface, contrasting with its typical increase toward the seabed on exterior walls. Although *M. senile* is highly tolerant to a large range of environmental conditions, including anoxia and a range of flow speeds and shows trophic plasticity (Mavraki, De Mesel, et al., 2020; Wahl, 1984), the present conditions may still affect them, as flow speeds have been shown to affect reproduction, morphology and prey capture (Anthony, 1997; Lesser et al., 1994). Amphipoda are sensitive to abiotic factors, such as DO, water movement, and sedimentation (Beermann & Franke, 2012; Conradi et al., 1997; Wu & Or, 2005). Jassa spp. generally show a preference for more exposed locations in the North Sea, potentially due to food availability and colonisation chances (Beermann & Franke, 2012). The reduced presence or competitive abilities of these species on the interior of the monopile, likely due to environmental stress, potentially alleviates competitive pressure, allowing other species to settle. However, some taxa observed with lower percentages on the outside, such as Serpulidae, may be equally present but obscured by overgrowth, leading to underrepresentation in video-based assessments. The co-occurrence of Ophiuroidea and Porifera corresponds to what is known from other locations and potentially reflects juvenile refuge formation or trophic facilitation (Oliveira et al., 2024; Turon et al., 2000).

Larval behaviour and dispersal further contribute to differences between interior and exterior communities. Jassa spp. lack planktonic larvae and rely on drifting individuals for dispersal (Havermans et al., 2007). Restricted water exchange may limit their colonisation success and explain their reduced presence on the interior walls. Many benthic organisms, including Amphipoda, *M. senile*, and Ascidiacea are capable of photoreception in their larval and adult stages and show behavioural responses to light stimuli (Kusakabe & Tsuda, 2007; Lynn et al., 2024; Navarro-Barranco & Hughes, 2015). *D. listerianum* larvae for instance prefer dim light just before settlement (Crisp & Ghobashy, 1971), meaning that individual behavioural patterns may affect settlement and survival success under these unique conditions, and thus contribute to the increased cover by Ascidiacea on the interior walls.

The interior conditions - restricted hydrodynamics, POC gradients, and absence of light - create conditions resembling marine caves, which are also characterised by distinct communities dominated by e.g., Porifera, Bryozoa, calcareous tube worms, or Ascidiacea (Gerovasileiou & Voultsiadou, 2012; Readman et al., 2024). Like marine caves, the monopile interiors exhibited small-scale patchiness (Denitto et al., 2007; Martí et al., 2004). The lower overall cover observed may correspond to slow colonisation, and community development known from marine caves (Denitto et al., 2007; Navarro-Barranco et al., 2015). This is likely driven by the limited flow and exchange of water, which strongly reduces the supply of settlement-competent larvae, combined with predation on these by the already established filter feeders (Palardy & Witman, 2014). While monopile interiors are structurally simpler than natural caves, these similarities suggest that they may function as analogous semi-enclosed habitats.

### 4.2. Variability between monopiles

Despite the clear patterns in some species and total cover, substantial variability in community composition was observed between monopiles, particularly on the interior walls. Variation on the interior walls may partly be explained by the differences in water quality due to the WRH configuration. However, because only F1 had a different WRH configuration, additional factors must contribute to the observed differences. Variation on the exterior walls was also evident, including the dominance of *M. gigas* on monopile F4. Small-scale environmental variations are likely contributors, as even limited distances can influence colonisation and community composition (Rule & Smith, 2005; Zintzen et al., 2008). Furthermore, the monopiles were installed in different seasons, with F1, E5, and D1 deployed in September and November, while F4 was installed in April of the following year. Variation in the timing of monopile placement may result in diverging community trajectories due to variability in larval supply and early recruitment processes, as well as differences in total cover (Underwood & Chapman, 2006).

### 4.3. Ecological risks and benefits

The observed patterns have important implications for potential benefits as well as risks associated with the implementation of WRHs. The development of a novel artificial habitat with distinct communities raises questions regarding its role in the stepping-stone effect and establishment of introduced non-indigenous species (iNIS ; Tyrrell & Byers, 2007). In this study, the iNIS *C. fornicata, M. gigas*, and several cryptic Ascidiacea, not identified up to species level but likely iNIS as well, were found on the exterior and interior of the monopile. However, *C. fornicata* and *M. gigas* exhibited a lower cover on the interior than on the exterior of the monopile, suggesting that the internal habitat is less suitable for these species. In contrast, the colonial Ascidiacea were more abundant on the interior, particularly in the shallow-to mid-depth zones, suggesting that the novel habitat may support specific iNIS that profit from reduced competition or are adapted to the internal environmental conditions. Taxonomic uncertainty limits the evaluation of ecological risks on the new habitat. Molecular approaches including eDNA, could help with early detection of iNIS.

The videos revealed notable accumulation of organic matter on the interior seafloor, suggesting reduced export under the limited water exchange. The observed grey mats are likely microbial mats, similar to those documented in marine environments with limited water exchange and often high organic inputs such as seasonally in Lake Grevelingen (The Netherlands), the Baltic Sea, marine caves, or the sediments underneath fish farms (Aranda et al., 2015; Gerovasileiou & Bianchi, 2021; Lipsewers et al., 2016; Noffke et al., 2016). The organic matter accumulation may increase microbial oxygen demand for degradation, which, together with the already lowered DO concentration near the seabed (Fig. 5), can result in hypoxic conditions at the sediment surface (Hargrave et al., 2008). Although the model already predicts decrease DO concentrations near the seabed, the real concentrations are likely lower than the predicted concentrations due to the biogeochemical processes occurring there. Persistent low-oxygen conditions can promote the development of anaerobic microbial communities and sulphur-oxidizing bacterial mats, while reducing macrofaunal abundance and altering benthic ecosystem processes (Conley et al., 2009; Grünke et al., 2011; Hargrave et al., 2008). A 16S rRNA gene analysis could identify the dominant taxa and better assess the ecological and biogeochemical implications of these communities (Lemoinne et al., 2024; van der Loos & Nijland, 2021). It is currently unclear, whether these conditions on the seafloor are seasonal or also remain during months with higher wave activity and consequent water exchange.

The installation of monopiles with WRHs approximately doubles the available vertical hard substrate in wind farms. Due to limited data on biomass, it remains unclear whether the biomass associated with interior communities represents a meaningful increase in total epifaunal biomass. This uncertainty limits possibilities to predict the magnitude of effects on biogeochemical processes and benthopelagic coupling.

The additional epifauna may also further enhance feeding opportunities and attract fish, while the interior could also function as shelter for species that utilise three dimensional or sheltered spaces as habitat. If optimised with the goal of not only reducing acidification but also to provide a meaningful increase of an asset of biodiversity of the wind farm, WRHs may function as a creative nature-inclusive design (Degraer et al., 2026) that enhance natural values beyond the natural local environment. Adjustments in number, orientation, or size may induce changes in habitat suitability and alter the epifaunal community as well as the potential for mobile species to use the interior as a shelter. The potential of using monopile interiors by fish and other mobile species remains unclear and warrants further investigation. Accessibility of the habitat to predatory species such as seals should also be considered.

The implementation of WRHs to prevent acidification on the interior of the monopiles may thus offer benefits and risks for the ecosystem. At present, the studied WRH configuration does not appear to support many iNIS or protected epifaunal species, although outcomes are likely to vary across locations due to differences in hydrodynamics, larvae availability, and overall habitat suitability. Importantly, the long-term community development and settlement of iNIS and protected species in this newly introduced habitat requires targeted ecological monitoring of the monopile interiors, which currently is not mandatory.

### 4.4. Limitations of the current study

The investigation of macrofauna was conducted using video recordings and percentage cover estimates, which limits taxonomic resolution and the detection of small or cryptic species. As a result, species-level patterns and the presence of inconspicuous taxa, including iNIS, are likely not reflected to their full extent. Nevertheless, we did find comparable numbers of taxonomic groups and phyla as other video-based studies on offshore production platforms, suggesting that broader community patterns were captured successfully (Schutter et al., 2019; Van der Stap et al., 2016). The sampling design was not fully balanced between interior and exterior recordings and monopiles, as the interior frames had a smaller AOI compared to the exterior and the number of analysed frames differed. Although the GLLVM accounted for these differences, especially rare, patchy, or small species might be affected by this difference. Differences in video quality might also generally affect the detectability of these species. Nevertheless, the main observed patterns are unlikely to be affected by this. Biomass estimation from video imagery proved challenging due to limited availability of appropriate reference data and the absence of laser lines on the interior transects, forcing an estimation of area assessed. While scrape samples collected by ROVs or divers would give a more accurate representation of community composition and biomass, such sampling was not feasible under the current operational and safety constraints. The biomass values were therefore only used as proxy input for the water quality model and not directly assessed to avoid overinterpretation.

The water quality modelling relied on several simplifying assumptions regarding biomass, ecophysiological parameters, and ambient conditions due to data and modelling limitations. Species-specific oxygen consumption and POC rates were not available for all taxa and were therefore approximated using literature values from a limited subset of species and then applied uniformly across monopiles, depth layers, and time. As a result, the modelling is not based on a true species-specific community. Biomass normalisation and conversion to carbon biomass using non-species-specific conversion factors introduced additional uncertainties. Furthermore, boundary conditions such as ambient oxygen and POC concentrations were held constant, thereby excluding natural temporal variability. This may obscure short-term or seasonal dynamics within the monopile. While the model provides valuable insight into how WRH configuration influences internal water exchange and identifies potential zones of limitation, it does not give information on absolute concentration levels, species-specific suitability, and fine scale vertical patterns. The modelling supports design sensitivity analysis and identification of limiting zones within the monopile, but does not provide species resolved ecological thresholds. It should therefore only be interpreted as a first-order description of biological demand rather than a detailed ecological budget.

## 5. Conclusion

This study demonstrates that offshore wind monopiles equipped with WRHs function as novel semi-enclosed artificial habitats that support epifaunal communities different from those on the exterior monopile surfaces. The combination of field observations and numerical modelling provides a unique insight into the relationship of the field-observed vertical structure of the epifaunal communities and the vertical gradients in DO and POC on the interior of the monopiles, generated by the WRHs.

Interior monopile surfaces were characterised by lower overall epifaunal cover, reduced dominance of typical offshore hard substrate taxa such as Amphipoda, Bivalvia, and Hexacorallia, and a comparatively heterogenous community including Ophiuroidea, Serpulidae, and Porifera. In addition, accumulation of organic matter and the presence of microbial mats on interior seafloors indicate altered breakdown of organic material and potential risks for seafloor epifauna.

Due to the absence of light, restricted hydrodynamics, and gradients in POC and DO, the monopile interiors do not simply resemble common hard substrate communities. Instead, they function as an artificial cave-like environment, that increases habitat heterogeneity within offshore wind farms and may contribute to local-scale biodiversity changes. However, ecological outcomes vary between monopiles and are influenced by structural configuration, highlighting the importance of design-specific effects. Together, these findings show how engineering modifications for improved corrosion protection also influence ecological functioning and provide a first step towards evaluating the ecological implications of WRH-equipped monopiles. Future work should focus on design improvements for a positive contribution to biodiversity assets and long-term abiotic and biotic monitoring, of species-level responses (including iNIS and protected species), habitat use by mobile species, and the microbial communities near the seabed.

## Acknowledgements

The authors gratefully acknowledge Vattenfall for data sharing, financial and operational support, and collaboration, in particular Iris Menger, Michail Karasoulas, and Tim Wilms. We thank the crew of Windcat Workboats for their support during offshore operations and Føn Energy Services for technical support and the deployment of the ROV inside and outside the monopile foundations. We also thank Oliver Bittner, Roos Demmers, Marjolein Kelder, Isaac de Boer-Ferrier, Oskar van Megen, and Francesca Di Paola for their assistance in data collection, processing and modelling. We acknowledge the contribution of NIOZ Royal Netherlands Institute for Sea Research, specifically Rob Witbaard, Furu Mienis, Yvo de Witte, and Dave Huijsman, for their support in the frame design and research setup and other valuable input. Additional thanks goes to Tjeerd Bouma and Ninon Mavraki for their contributions to the project development. This work was carried out in collaboration with partners within the JIP-LIFE project, including Wageningen Marine Research, Deltares, Waardenburg Ecology, Seaward, and The Rich North Sea programme. The research was funded by TKI Offshore Energy and Netherlands Enterprise Agency (RVO) under the JIP-LIFE project (grant number TKITOE_WOZ_2410, 2025). The Dutch Postcode Lottery funded The Rich North Sea programme as a Dream Fund project executed by The North Sea Foundation and Natuur & Milieu. Part of the work by Joop Coolen was funded through the NWO-NWA-ORC funded NO-REGRETS project (grant ID NWA.1630.23.016). Leandra Kornau was funded by TKI Deltatechnology and INSITE North Sea via the ASSESS project with co-funding from Heerema Marine Contractors.

## Declaration of generative AI and AI-assisted technologies in the manuscript preparation process

During the preparation of this work the author(s) used Microsoft Copilot and OpenAI ChatGPT to assist in language polishing. After using this tool/service, the authors reviewed and edited the content as needed and take full responsibility for the content of the published article.

## Supplementary

### Supplementary Data 1: Method Details

### Supplementary Data 2: Abiotic data collected at F1

Table Supplementary Data 2

### Supplementary Data 3: Raw and averaged percentage cover data

Table Supplementary Data 3

### Supplementary Data 4: GLLVM model summary data

Table Supplementary Data 4

## References

Akaike, H. (1973). Information Theory and an Extension of the Maximum Likelihood Principle. International Symposium on Information Theory, 267–281.

Andersen, L. D. (2017). Renewal of water in monopiles for offshore wind [Aalborg University]. http://www.ses.aau.dk/studienaevn/byggeri/ (accessed 03 June 2026).

Anthony, K. R. N. (1997). Prey Capture by the Sea Anemone Metridium senile (L.): Effects of Body Size, Flow Regime, and Upstream Neighbors. The Biological Bulletin, 192(1), 73–86. 10.2307/1542577

Aranda, C. P., Valenzuela, C., Matamala, Y., Godoy, F. A., & Aranda, N. (2015). Sulphur-cycling bacteria and ciliated protozoans in a Beggiatoaceae mat covering organically enriched sediments beneath a salmon farm in a southern Chilean fjord. Marine Pollution Bulletin, 100(1), 270–278. 10.1016/J.MARPOLBUL.2015.08.040

Beermann, J. (2013). Ecological differentiation among amphipod species in marine fouling communities [FU]. 10.17169/REFUBIUM-9982

Beermann, J., & Franke, H. D. (2012). Differences in resource utilization and behaviour between coexisting Jassa species (Crustacea, Amphipoda). Marine Biology, 159(5), 951–957. 10.1007/S00227-011-1872-7

Carstensen, S. (2025). Preprint: A mathematical model for the replenishment of water inside Monopile. https://www.researchgate.net/publication/388552572_A_mathematical_model_for_the_replenishment_of_water_inside_Monopile (accessed 03 June 2026).

Conley, D. J., Björck, S., Bonsdorff, E., Carstensen, J., Destouni, G., Gustafsson, B. G., Hietanen, S., Kortekaas, M., Kuosa, H., Meier, H. E. M., Müller-Karulis, B., Nordberg, K., Norkko, A., Nürnberg, G., Pitkänen, H., Rabalais, N. N., Rosenberg, R., Savchuk, O. P., Slomp, C. P., … Zillén, L. (2009). Hypoxia-Related Processes in the Baltic Sea. Environmental Science and Technology, 43(10), 3412–3420. 10.1021/ES802762A

Connell, J. H., & Slatyer, R. O. (1977). Mechanisms of Succession in Natural Communities and Their Role in Community Stability and Organization. The American Naturalist, 111(982), 1119–1144.

Conradi, M., López-González, P. J., & García-Gomez, C. (1997). The amphipod community as a bioindicator in Algeciras Bay (Southern Iberian Peninsula) based on a spatio-temporal distribution. Marine Ecology, 18(2), 97–111. 10.1111/j.1439-0485.1997.tb00430.x

Coolen, J. W. P., Bittner, O., Driessen, F. M. F., Van Dongen, U., Siahaya, M. S., De Groot, W., Mavraki, N., Bolam, S. G., & Van der Weide, B. (2020). Ecological implications of removing a concrete gas platform in the North Sea. Journal of Sea Research, 166, 101968. 10.1016/J.SEARES.2020.101968

Coolen, J. W. P., Van der Weide, B., Bittner, O., Peck, N., Kornau, L., Keur, M., Foekema, E., Ibanez-Erquiaga, B., & Huizinga, J.-J. (2025). Marine growth sampling tool evaluationrJ: evaluation of the performance of an ROV-mounted tool for sampling marine growth on Offshore energy structures. 10.18174/678892

Coolen, J. W. P., Van Der Weide, B., Cuperus, J., Blomberg, M., Van Moorsel, G. W. N. M., Faasse, M. A., Bos, O. G., Degraer, S., & Lindeboom, H. J. (2020). Benthic biodiversity on old platforms, young wind farms, and rocky reefs. ICES Journal of Marine Science, 77(3), 1250–1265. 10.1093/ICESJMS/FSY092

Coolen, J. W. P., Vanaverbeke, J., Dannheim, J., Garcia, C., Birchenough, S. N. R., Krone, R., & Beermann, J. (2022). Generalized changes of benthic communities after construction of wind farms in the southern North Sea. Journal of Environmental Management, 315, 115173. 10.1016/J.JENVMAN.2022.115173

Crisp, D. J., & Ghobashy, A. F. A. A. (1971). Responses of the larva of Diplosoma listerianum to light and gravity. In D. J. Crisp (Ed.), Fourth European Marine Biology Symposium (pp. 443–465). Cambridge University Press.

Dannheim, J., Beermann, J., Coolen, J. W. P., Vanaverbeke, J., Degraer, S., Birchenough, S. N. R., Garcia, C., Lacroix, G., Fiorentino, D., Lindeboom, H., Krone, R., Pehlke, H., Braeckman, U., & Brey, T. (2025). Offshore wind turbines constitute benthic secondary production hotspots on and around constructions. Journal of Environmental Management, 393, 126922. 10.1016/J.JENVMAN.2025.126922

Dannheim, J., Kloss, P., Vanaverbeke, J., Mavraki, N., Zupan, M., Spielmann, V., Degraer, S., Birchenough, S. N. R., Janas, U., Sheehan, E., Teschke, K., Gill, A. B., Hutchison, Z., Carey, D. A., Rasser, M., Buyse, J., Van der Weide, B., Bittner, O., Causon, P., … Coolen, J. W. P. (2025). Biodiversity Information of benthic Species at ARtificial structures – BISAR. Scientific Data 2025 12:1, 12(1), 1–11. 10.1038/s41597-025-04920-1

De Borger, E., Ivanov, E., Capet, A., Braeckman, U., Vanaverbeke, J., Grégoire, M., & Soetaert, K. (2021). Offshore Windfarm Footprint of Sediment Organic Matter Mineralization Processes. Frontiers in Marine Science, 8, 632243. 10.3389/FMARS.2021.632243

Degraer, S., Carey, D. A., Coolen, J. W. P., Hutchison, Z. L., Kerckhof, F., Rumes, B., & Vanaverbeke, J. (2020). Offshore wind farm artificial reefs affect ecosystem structure and functioning: A synthesis. Oceanography, 33(4), 48–57. 10.5670/OCEANOG.2020.405

Degraer, S., Cornacchia, L., Ziemba, A., Agardy, T., Van den Burg, S. W. K., Van de Braak, K., Van Duren, L. A., Declercq, A., Eggermont, M., Van Gerven, A., Kerkhove, T. R. H., Moulaert, I., Patel, T., Petersen, J. K., Poelman, M., Stratigaki, V., Strothotte, E., Vanaverbeke, J., & El Serafy, G. (2026). Rethinking the responsible application of nature-inclusive design in marine infrastructure to restore ecosystems and enhance biodiversity. ICES Journal of Marine Science, 83(4). 10.1093/ICESJMS/FSAG050

Deltares. (2022). D-Flow Flexible Mesh: Computational Core and user Interface - User Manual.

Denitto, F., Terlizzi, A., & Belmonte, G. (2007). Settlement and primary succession in a shallow submarine cave: Spatial and temporal benthic assemblage distinctness. Marine Ecology, 28(S1), 35–46. 10.1111/j.1439-0485.2007.00172.x

Fugro Survey B.V. (2016). Site Studies Wind Farm Zone Hollandse Kust (zuid). https://offshorewind.rvo.nl/file/download/3a901fda-60ec-4fc8-8c27-a8a4f5207d86/1483696372hkz_20160824_fugro_geophysical%20report_wfs%20i_gh176-r1_revb_f_compleet_v2.pdf

Gerovasileiou, V., & Bianchi, C. N. (2021). Mediterranean marine caves: A synthesis of current knowledge. In S. J. Hawkins, A. J. Lemasson, A. L. Allock, A. E. Bates, M. Byrne, A. J. Evans, L. B. Firth, E. M. Marzinelli, B. D. Russel, I. P. Smith, S. E. Swearer, & P. A. Todd (Eds.), Oceanography and Marine Biology: An Annual Review, Volume 59 (1st ed., Vol. 59, pp. 1–88). CRC Press EBooks. 10.1201/9781003138846-1

Gerovasileiou, V., & Voultsiadou, E. (2012). Marine Caves of the Mediterranean Sea: A Sponge Biodiversity Reservoir within a Biodiversity Hotspot. PLOS ONE, 7(7), e39873. 10.1371/JOURNAL.PONE.0039873

Glarou, M., Zrust, M., & Svendsen, J. C. (2020). Using Artificial-Reef Knowledge to Enhance the Ecological Function of Offshore Wind Turbine Foundations: Implications for Fish Abundance and Diversity. Journal of Marine Science and Engineering 2020, Vol. 8, Page 332, 8(5), 332. 10.3390/JMSE8050332

[dataset]Global Administrative Areas. (2025). GADM database of Global Administrative Areas, version 4.1. University of Berkeley, Museum of Vertebrate Zoology and the International Rice Research Institute (accessed 26 May 2026).

Grünke, S., Felden, J., Lichtschlag, A., Girnth, A. C., De Beer, D., Wenzhöfer, F., & Boetius, A. (2011). Niche differentiation among mat-forming, sulfide-oxidizing bacteria at cold seeps of the Nile Deep Sea Fan (Eastern Mediterranean Sea). Geobiology, 9(4), 330–348. 10.1111/j.1472-4669.2011.00281.x

Hargrave, B. T., Holmer, M., & Newcombe, C. P. (2008). Towards a classification of organic enrichment in marine sediments based on biogeochemical indicators. Marine Pollution Bulletin, 56(5). 10.1016/j.marpolbul.2008.02.006

Harrington, C. T. (2010). The invasion of the colonial ascidian Didemnum vexillum, into south shore bays of Long Island, New York-Feeding and metabolic characteristics [Stony Brook University]. In Stony Brook Theses and Dissertations Collection, 2006-2020 (closed to submissions). https://commons.library.stonybrook.edu/stony-brook-theses-and-dissertations-collection/92/

Havermans, C., De Broyer, C., Mallefet, J., & Zintzen, V. (2007). Dispersal mechanisms in amphipods: A case study of Jassa herdmani (Crustacea, Amphipoda) in the North Sea. Marine Biology, 153(1), 83–89. 10.1007/S00227-007-0788-8

Hilbert, L. R., Black, A. R., Andersen, F., & Mathiesen, T. (2011). Inspection and monitoring of corrosion inside monopile foundations for offshore wind turbines (EUROCORR 2011 Conference).

Jacobsen, N. G., Hoballah Jalloul, M., Welzel, M., Schendel, A., & Carstensen, S. (2025). Jet velocities in water replenishment holes: A large-scale experiment. Ocean Engineering, 342, 122867. 10.1016/J.OCEANENG.2025.122867

Kesgin, E., Christensen Sørensen, J., Jacobsen, N. G., & Carstensen, S. (2024). Experimental Study on Water Replenishment Holes in Offshore Monopiles. Ocean Engineering, 306. 10.1016/j.oceaneng.2024.118071

Kindeberg, T., Attard, K. M., Hüller, J., Müller, J., Quintana, C. O., & Infantes, E. (2024). Structural complexity and benthic metabolism: resolving the links between carbon cycling and biodiversity in restored seagrass meadows. Biogeosciences, 21(7), 1685–1705. 10.5194/BG-21-1685-2024

Korhonen, P. (2025). Analysing sparse ecological percent cover data using gllvm. https://cran.r-project.org/web/packages/gllvm/vignettes/vignette8.html (accessed 03 June 2026).

Korhonen, P., Hui, F. K. C., Niku, J., & Taskinen, S. (2023). Fast and universal estimation of latent variable models using extended variational approximations. Statistics and Computing, 33(1). 10.1007/S11222-022-10189-W

Korhonen, P., Hui, F. K. C., Niku, J., Taskinen, S., & Van der Veen, B. (2024). A comparison of joint species distribution models for percent cover data. Methods in Ecology and Evolution, 15(12), 2359–2372. 10.1111/2041-210X.14437

Krone, R., Gutow, L., Joschko, T. J., & Schröder, A. (2013). Epifauna dynamics at an offshore foundation – Implications of future wind power farming in the North Sea. Marine Environmental Research, 85, 1–12. 10.1016/J.MARENVRES.2012.12.004

Kumala, L., Thomsen, M., & Canfield, D. E. (2023). Respiration kinetics and allometric scaling in the demosponge Halichondria panicea. BMC Ecology and Evolution 2023 23:1, 23(1), 53-. 10.1186/S12862-023-02163-5

Kusakabe, T., & Tsuda, M. (2007). Photoreceptive Systems in Ascidians. Photochemistry and Photobiology, 83(2), 248–252. 10.1562/2006-07-11-IR-965

Lemoinne, A., Dirberg, G., Georges, M., & Robinet, T. (2024). Evaluation of a Nanopore Sequencing Strategy on Bacterial Communities From Marine Sediments. Environmental DNA, 6(5), e70009. 10.1002/EDN3.70009

Lesser, M. P., Witman, J. D., & Sebens, K. P. (1994). Effects of Flow and Seston Availability on Scope for Growth of Benthic Suspension-Feeding Invertebrates from the Gulf of Maine. The Biological Bulletin, 187(3), 319–335. 10.2307/1542289

Lipsewers, Y. A., Hopmans, E. C., Meysman, F. J. R., Sinninghe Damsté, J. S., & Villanueva, L. (2016). Abundance and diversity of denitrifying and anammox bacteria in seasonally hypoxic and sulfidic sediments of the saline lake Grevelingen. Frontiers in Microbiology, 7(OCT). 10.3389/FMICB.2016.01661

Lynn, K. D., Quintanilla-Ahumada, D., Duarte, C., & Quijón, P. A. (2024). Artificial light at night alters the feeding activity and two molecular indicators in the plumose sea anemone Metridium senile (L.). Marine Pollution Bulletin, 202, 116352. 10.1016/J.MARPOLBUL.2024.116352

Mackinson, S., & Daskalov, G. (2007). An ecosystem model of the North Sea to support an ecosystem approach to fisheries management: description and parameterisation. Science Series Technical Report no. 142.

Maher, M. M. (2018). The Corrosion and Biofouling Characteristics of Sealed vs. Perforated Offshore Monopile Interiors: Experiment Design Comparing Corrosion and Environment Inside Steel Pipe [Florida Institute of Technology ]. In Theses and Dissertations. https://repository.fit.edu/etd/1209

Maher, M. M., & Swain, G. (2018). The Corrosion and Biofouling Characteristics of Sealed vs. Perforated Offshore Monopile Interiors Experiment Design Comparing Corrosion and Environment Inside Steel Pipe. Proceedings of OCEANS 2018 MTS/IEEE Charleston, 22–25. 10.1109/OCEANS.2018.8604522

Martí, R., Uriz, M. J., Ballesteros, E., & Turon, X. (2004). Temporal variation of several structure descriptors in animal-dominated benthic communities in two Mediterranean caves. Journal of the Marine Biological Association of the United Kingdom, 84(3), 573–580. 10.1017/S0025315404009579H

Mavraki, N., De Mesel, I., Degraer, S., Moens, T., & Vanaverbeke, J. (2020). Resource Niches of Co-occurring Invertebrate Species at an Offshore Wind Turbine Indicate a Substantial Degree of Trophic Plasticity. Frontiers in Marine Science, 7, 538083. 10.3389/FMARS.2020.00379

Mavraki, N., Degraer, S., Moens, T., & Vanaverbeke, J. (2020). Functional differences in trophic structure of offshore wind farm communities: A stable isotope study. Marine Environmental Research, 157, 104868. 10.1016/J.MARENVRES.2019.104868

Mavraki, N., Degraer, S., Vanaverbeke, J., & Braeckman, U. (2020). Organic matter assimilation by hard substrate fauna in an offshore wind farm area: a pulse-chase study. ICES Journal of Marine Science, 77(7–8), 2681–2693. 10.1093/ICESJMS/FSAA133

Migné, A., & Davoult, D. (1997). Oxygen consumption in two benthic cnidarians: Alcyonium digitatum (Linnaeus, 1758) and Urticina felina (Linnaeus, 1767). Proc. 6th Int. Conf. on Coelenterate Biology, 1995, 321–328.

Navarro-Barranco, C., Guerra-García, J. M., Sánchez-Tocino, L., Ros, M., Florido, M., & García-Gómez, J. C. (2015). Colonization and successional patterns of the mobile epifaunal community along an environmental gradient in a marine cave. Marine Ecology Progress Series, 521, 105–115. 10.3354/MEPS11126

Navarro-Barranco, C., & Hughes, L. E. (2015). Effects of light pollution on the emergent fauna of shallow marine ecosystems: Amphipods as a case study. Marine Pollution Bulletin, 94(1–2), 235–240. 10.1016/j.marpolbul.2015.02.023

Nelson, M. L., & Craig, S. F. (2011). Role of the sea anemone Metridium senile in structuring a developing subtidal fouling community. Marine Ecology Progress Series, 421, 139–149. 10.3354/MEPS08838

Niku, J., Hui, F. K. C., Taskinen, S., & Warton, D. I. (2019). gllvm: Fast analysis of multivariate abundance data with generalized linear latent variable models in r. Methods in Ecology and Evolution, 10(12), 2173–2182. 10.1111/2041-210X.13303

Noffke, A., Sommer, S., Dale, A. W., Hall, P. O. J., & Pfannkuche, O. (2016). Benthic nutrient fluxes in the Eastern Gotland Basin (Baltic Sea) with particular focus on microbial mat ecosystems. Journal of Marine Systems, 158, 1–12. 10.1016/J.JMARSYS.2016.01.007

Noisette, F., Bordeyne, F., Davoult, D., & Martin, S. (2016). Assessing the physiological responses of the gastropod Crepidula fornicata to predicted ocean acidification and warming. Limnology and Oceanography, 61(2), 430–444. 10.1002/LNO.10225

Oliveira, M., da Silva, M. Q. M., Barroso, C. X., Salani, S., & Matthews-Cascon, H. (2024). A report on new sponge-ophiuroid associations and reinforcement of scientific knowledge. Ocean and Coastal Research, 72, e24007. 10.1590/2675-2824072.22152

Ortega, M. M., Lopez de Pariza, J. M., & Navarro, E. (1988). Seasonal changes in the biochemical composition and oxygen consumption of the sea anemone Actinia equina as related to body size and shore level. Marine Biology, 97(1), 137–143. 10.1007/BF00391253

Pack, K. E., Rius, M., & Mieszkowska, N. (2021). Long-term environmental tolerance of the non-indigenous Pacific oyster to expected contemporary climate change conditions. Marine Environmental Research, 164, 105226. 10.1016/J.MARENVRES.2020.105226

Palardy, J. E., & Witman, J. D. (2014). Flow, recruitment limitation, and the maintenance of diversity in marine benthic communities. Ecology, 95(2), 286–297. 10.1890/12-1612.1

Posit Team. (2025). RStudio: Integrated Development Environment for R. Posit Software, PBC.

Postma, L., Boderie, P. M. A., Van Gils, J. A. G., & Van Beek, J. K. L. (2003). Component Software Systems for Surface Water Simulation. Lecture Notes in Computer Science (Including Subseries Lecture Notes in Artificial Intelligence and Lecture Notes in Bioinformatics), 2657, 649–658. 10.1007/3-540-44860-8_67

R Core Team. (2025). R: A Language and Environment for Statistical Computing (V.4.5.0). R Foundation for Statistical Computing.

Readman, J. A. J., Tyler-Walters, H., & Watson, A. J. (2024). Sponges, bryozoans and ascidians on deeply overhanging lower shore bedrock or caves - MarLIN - The Marine Life Information Network. Marine Life Information Network: Biology and Sensitivity Key Information Reviews. https://www.marlin.ac.uk/habitats/detail/358/sponges_bryozoans_and_ascidians_on_deeply_overhanging_lower_shore_bedrock_or_caves

Ricciardi, A., & Bourget, E. (1998). Weight-to-weight conversion factors for marine benthic macroinvertebrates. Marine Ecology Progress Series, 163, 245–251. 10.3354/MEPS163245

Riisgård, H. U., & Larsen, P. S. (2015). Filter-Feeding Zoobenthos and Hydrodynamics. In S. Rossi, L. Bramanti, A. Gori, & C. Orejas (Eds.), Marine Animal Forests (pp. 1–25). Springer, Cham. 10.1007/978-3-319-17001-5_19-1

Rijksoverheid. (2022). Shell en Eneco winnen tender windpark op zee Hollandse Kust (west). https://www.rijksoverheid.nl/actueel/nieuws/2022/12/15/shell-en-eneco-winnen-tender-windpark-op-zee-hollandse-kust-west (accessed 16 August 2023).

[dataset]Rijkswaterstaat. (2024). Bathymetry Dutch part of the North Sea deeper than 10 m LAT. http://Data.Europa.Eu/88u/Dataset/A322184d-6285-4856-Bb78-F450b30ffc0e (accessed 26 May 2026).

[dataset]Rijkswaterstaat, & Deltares. (2024). Positions of turbines and licensed wind farm areas (Netherlands). https://Viewer.Openearth.Nl/Ihm-Viewer/Download/Geoserver?Layers=85222896 (accessed 26 May 2026).

Rueda, J. L., Smaal, A. C., & Scholten, H. (2005). A growth model of the cockle (Cerastoderma edule L.) tested in the Oosterschelde estuary (The Netherlands). Journal of Sea Research, 54(4), 276–298. 10.1016/J.SEARES.2005.06.001

Rule, M. J., & Smith, S. D. A. (2005). Spatial variation in the recruitment of benthic assemblages to artificial substrata. Marine Ecology Progress Series, 290, 67–78. 10.3354/MEPS290067

Sarada, S., Xie, S., Gupta, A., & Desai, A. (2025). CFD Analysis of Pressure Variations on Airtight Platforms for Offshore Wind Turbines Monopile Foundation. Proceedings of the Annual Offshore Technology Conference. 10.4043/35970-MS

Schutter, M., Dorenbosch, M., Driessen, F. M. F., Lengkeek, W., Bos, O. G., & Coolen, J. W. P. (2019). Oil and gas platforms as artificial substrates for epibenthic North Sea fauna: Effects of location and depth. Journal of Sea Research, 153, 101782. 10.1016/J.SEARES.2019.101782

Souster, T. A., Morley, S. A., & Peck, L. S. (2018). Seasonality of oxygen consumption in five common Antarctic benthic marine invertebrates. Polar Biology 2018 41:5, 41(5), 897–908. 10.1007/S00300-018-2251-3

ter Hofstede, R., Bouma, T. J., & Van Koningsveld, M. (2023). Five golden principles to advance marine reef restoration by linking science and industry. Frontiers in Marine Science, 10, 1143242. 10.3389/FMARS.2023.1143242

Tupkar, S. A., & Sappe Narasimhamurthy, S. (2022). CFD Analysis of Water Replenishment Holes in an Offshore Wind Turbine Foundation [Linköping University]. https://urn.kb.se/resolve?urn=urn:nbn:se:liu:diva-182977 (accessed 04 May 2026).

Turon, X., Codina, M., Tarjuelo, I., Uriz, M. J., & Becerro, M. A. (2000). Mass recruitment of Ophiothrix fragilis (Ophiuroidea) on sponges: settlement patterns and post-settlement dynamics. Marine Ecology Progress Series, 200, 201–212. 10.3354/MEPS200201

Tyrrell, M. C., & Byers, J. E. (2007). Do artificial substrates favor nonindigenous fouling species over native species? Journal of Experimental Marine Biology and Ecology, 342, 54–60. 10.1016/J.JEMBE.2006.10.014

Underwood, A. J., & Chapman, M. G. (2006). Early development of subtidal macrofaunal assemblages: relationships to period and timing of colonization. Journal of Experimental Marine Biology and Ecology, 330(1), 221–233. 10.1016/J.JEMBE.2005.12.029

van der Loos, L. M., & Nijland, R. (2021). Biases in bulk: DNA metabarcoding of marine communities and the methodology involved. Molecular Ecology, 30(13), 3270–3288. 10.1111/MEC.15592

Van der Stap, T., Coolen, J. W. P., & Lindeboom, H. J. (2016). Marine Fouling Assemblages on Offshore Gas Platforms in the Southern North Sea: Effects of Depth and Distance from Shore on Biodiversity. PLOS ONE, 11(1), e0146324. 10.1371/JOURNAL.PONE.0146324

Van der Veen, B. (2025). GLLVM-workshop: Physialia workshop on Generalized Linear Latent Variable Models. https://github.com/BertvanderVeen/GLLVM-workshop (accessed 26 May 2026).

Van Duren, L. A., Zijl, F., Van Kessel, T., Van Zeist, V. T. M., Vilmin, L. M., Van der Meer, J., Aarts, G., Van der Molen, J., Soetaert, K., & Minns, A. M. (2021). Ecosystem effects of large upscaling of offshore wind on the North Sea – Synthesis report.

Van Sluis, C. J., Van Onselen, E., Airoldi, L., Duarte, C. M., Van Rijswick, H. F. M. W., Van der Heide, T., Olie, R. A., Kelder, M., & Bouma, T. J. (2025). Financing marine restoration through offshore wind investments. BioScience, 75(10), 856–864. 10.1093/BIOSCI/BIAF092

Voet, H. E. E., Van Colen, C., & Vanaverbeke, J. (2022). Climate change effects on the ecophysiology and ecological functioning of an offshore wind farm artificial hard substrate community. Science of The Total Environment, 810, 152194. 10.1016/J.SCITOTENV.2021.152194

Wahl, M. (1984). The fluffy sea anemone Metridium senile in periodically oxygen depleted surroundings. Marine Biology, 81(1), 81–86. 10.1007/BF00397629

Walker, S. J., Schlacher, T. A., & Schlacher-Hoenlinger, M. A. (2007). Spatial heterogeneity of epibenthos on artificial reefs: Fouling communities in the early stages of colonization on an East Australian shipwreck. Marine Ecology, 28(4), 435–445. 10.1111/J.1439-0485.2007.00193

Wickham, H. (2016). ggplot2. Springer International Publishing. 10.1007/978-3-319-24277-4

Wijsman, J. W. M., Herman, P. M. J., & Gomoiu, M. T. (1999). Spatial distribution in sediment characteristics and benthic activity on the northwestern Black Sea shelf. Marine Ecology Progress Series, 181, 25–39. 10.3354/MEPS181025

Wildish, D., & Kristmanson, D. (1997). Benthic Suspension Feeders and Flow. Benthic Suspension Feeders and Flow. 10.1017/CBO9780511529894

World Economic Forum. (2025). Nature Positive: Role of the Offshore Wind Sector. https://reports.weforum.org/docs/WEF_Nature_Positive_Role_of_the_Offshore_Wind_Sector.pdf (accessed 31 May 2026).

Wu, R. S. S., & Or, Y. Y. (2005). Bioenergetics, growth and reproduction of amphipods are affected by moderately low oxygen regimes. Marine Ecology Progress Series, 297, 215–223. 10.3354/MEPS297215

Zintzen, V., Norro, A., Massin, C., & Mallefet, J. (2008). Spatial variability of epifaunal communities from artificial habitat: Shipwrecks in the Southern Bight of the North Sea. Estuarine, Coastal and Shelf Science, 76(2), 327–344. 10.1016/J.ECSS.2007.07.012

Zupan, M., Coolen, J., Mavraki, N., Degraer, S., Moens, T., Kerckhof, F., Lopez Lopez, L., & Vanaverbeke, J. (2024). Life on every stone: Characterizing benthic communities from scour protection layers of offshore wind farms in the southern North Sea. Journal of Sea Research, 201, 102522. 10.1016/J.SEARES.2024.102522

Zupan, M., Rumes, B., Vanaverbeke, J., Degraer, S., & Kerckhof, F. (2023). Long-Term Succession on Offshore Wind Farms and the Role of Species Interactions. Diversity, 15(2), 288. 10.3390/D15020288/S1

Zuur, A. F., & Ieno, E. N. (2025). The world of zero-inflated models. Volume 3: Using GLLVM. Highland Statistics Ltd.

