## Supplementary Data 1 for "Renewable energy caves: water replenishment holes in offshore monopiles create novel marine habitats"

### Supplementary Data 1: Method Details

#### 1.1 Tables and Figures

Table S1. Technical background of investigated monopiles.

| Monopile | Water depth (mLAT) | Lower WRH orientation (°N) | Lower WRH elevation (mLAT) | Upper WRH orientation (°N) | SPL installation date | MP installation date |
|---|---|---|---|---|---|---|
| **E5** | -25.24 | 191, 258 | -20.89 | 191, 255 | 1 July 2021 | 3 November 2021 |
| **D1** | -23.26 | 191, 258 | -18.91 | 191, 255 | 24 June 2021 | 1 September 2021 |
| **F1** | -24.53 | 191, 25 | -20.18 | 191, 255 | 14 August 2021 | 8 September 2021 |
| **F4** | -24.97 | 191, 258 | -20.62 | 191, 255 | 1 July 2021 | 21 April 2022 |

Table S2. Monitoring dates for abiotic and biotic monitoring data used in this study, collected around the four monopiles.

| Monopile | Abiotic (Probes) | Biotic (Video) | |
|---|---|---|---|
| | | *Exterior* | *Interior* |
| **E5** | - | 2 September 2024 | 9 September 2024 |
| **D1** | - | 2 September 2024 | 5 September 2024 |
| **F1** | 16 October 2024 | 2 September 2024 | 6 September 2024 |
| **F4** | - | 30 August 2024 | 7 September 2024 |

Table S3. Number of analysed transects stills (frames) from the exterior and interior of the four monopiles.

| **Position/MP** | | **D1** | **E5** | **F1** | **F4** |
|---|---|---|---|---|---|
| **Transects** | Interior | 4 | 4 | 4 | 4 |
| | Exterior | 2 | 2 | 2 | 3 |
| **Frames** | Interior | 77 | 87 | 53 | 100 |
| | Exterior | 40 | 43 | 30 | 44 |

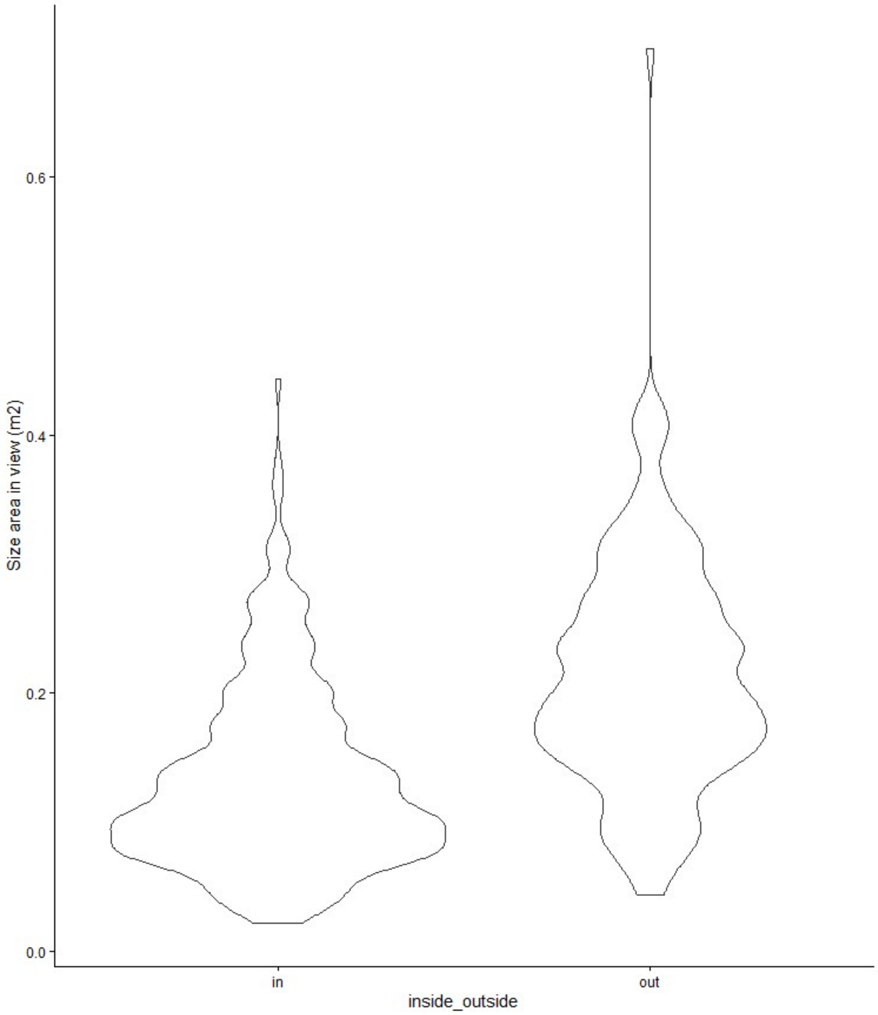


Fig. S1: Area of interest (AOI) in $m^2$ assessed for the different frames inside versus outside.

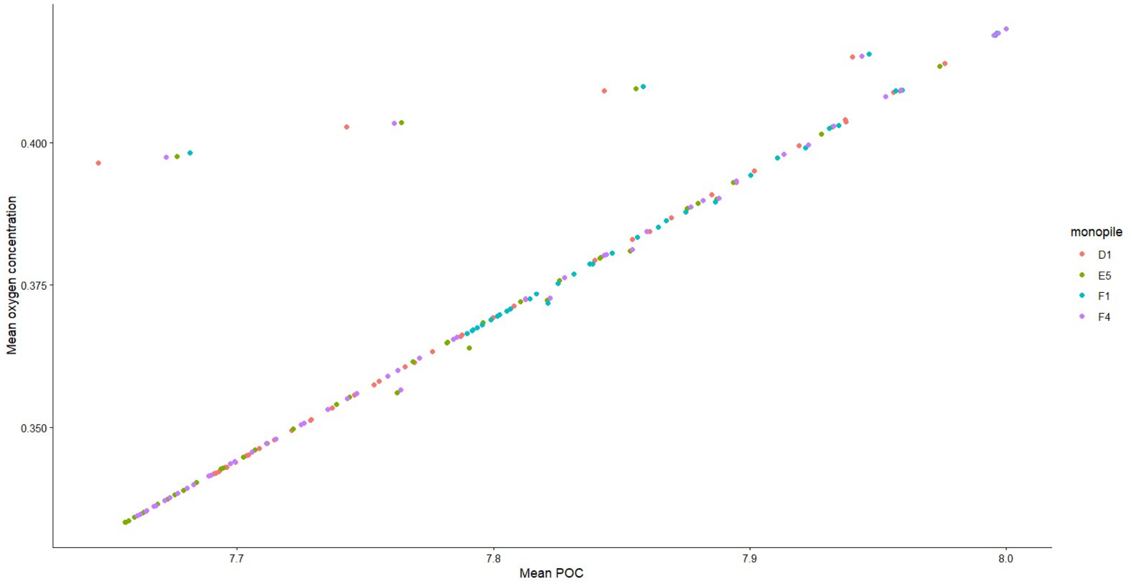


Fig. S2: Oxygen concentration vs POC showed strong collinearity.

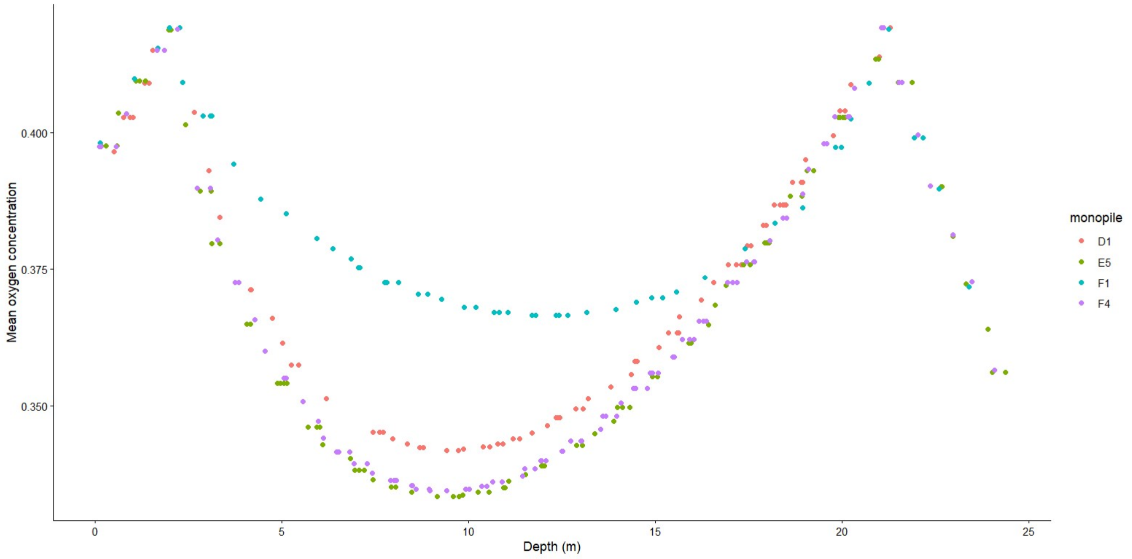


Fig. S3: Depth and oxygen concentration showed strong non-linear collinearity.

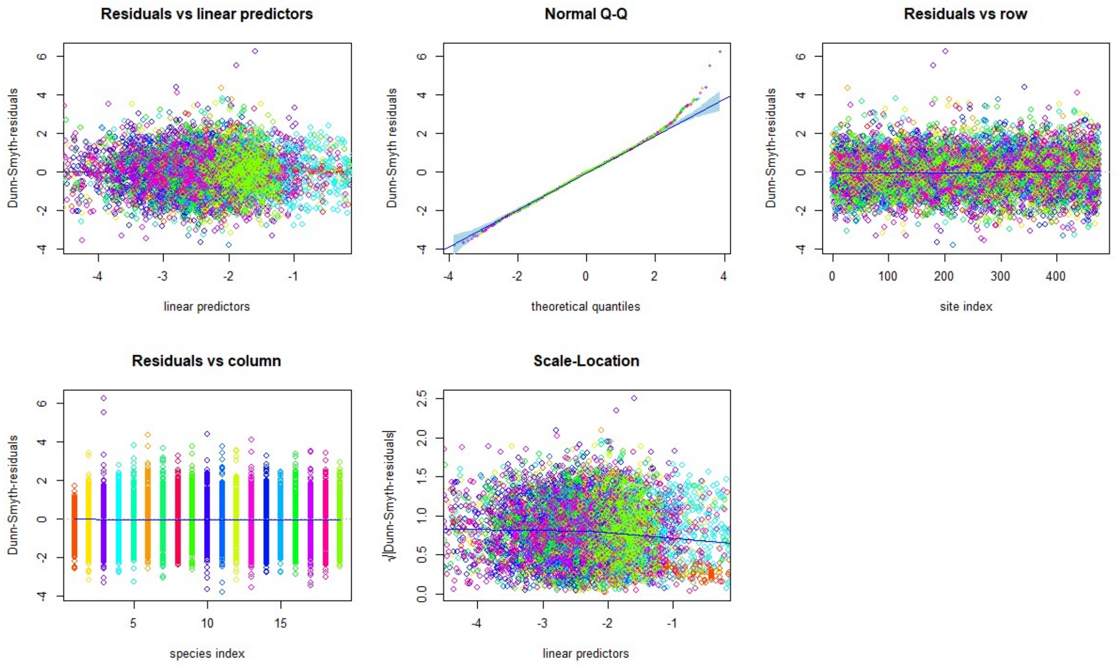


Fig. S4: GLLVM model validation plots.

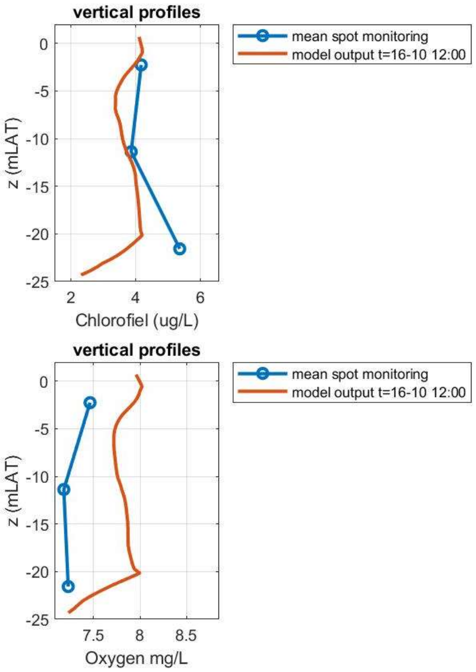


Fig. S5: Measured (blue) and modelled (orange) dissolved oxygen and chlorophyll concentrations (chlorophyll measured; chlorophyll modelled via POC1 conversion) during the monitoring campaign on 16 October 2024.

#### 1.2 Detailed coupled hydrodynamic and water quality model description

*Hydrodynamic water replenishment model*

Water exchange between the ambient sea and the monopile interior was simulated using a 1DV model, representing the monopile as a vertical water column and a compressible air pocket beneath the airtight platform. Exchange with the surrounding sea occurs through WRHs at prescribed elevations, while horizontal variability is neglected and cross-sections are assumed well mixed.

Inflow and outflow through the WRHs are driven by time-varying pressure differences resulting from water-level fluctuations, wave-induced dynamic pressures, and current-induced pressures around the monopile circumference. Air trapped above the internal water column can escape through ventilation openings, while compression and expansion of the enclosed air volume provide additional damping of internal water-level oscillations. The flow through each WRH is computed using an unsteady Bernoulli formulation with hydraulic loss coefficients representing contraction, expansion, and jet losses.

The model was applied to monopiles at the Hollandse Kust Zuid offshore wind farm using as-built geometries and WRH configurations. Hourly metocean forcing was prescribed using spectral wave parameters from a regional SWAN model and water levels and depth-averaged currents from the Dutch Continental Shelf Model (Deltares, 2022). Wave-induced pressures were derived using linear wave theory applied to stochastic JONSWAP spectra, with water-level and current-induced pressures added through linear superposition. The resulting exchange fluxes form the hydrodynamic input to the water quality model.

The hydrodynamic replenishment model was verified against laboratory wave experiments and calibrated loss coefficients, reproducing internal water-level oscillations and exchange fluxes within approximately 10 %, indicating that the dominant replenishment processes are adequately represented.

*Water quality model*

Internal water quality was simulated using the open-source DELWAQ model (Postma et al., 2003) in a 1DV configuration. The monopile interior was discretised into 61 vertical layers with variable thickness reflecting the tapered geometry. Horizontal gradients were neglected. Exchange fluxes from the hydrodynamic model were imposed as advective transports between layers. Layers above the upper WRH were allowed to adjust dynamically with internal water-level variations, while layers below the lower WRH experienced recirculation driven by inflow and outflow at that elevation.

Two state variables were modelled as primary indicators of habitat suitability: DO and POC, the latter acting as a proxy for food availability. The model includes vertical advection, vertical diffusion, sedimentation of POC, microbial decomposition of organic matter, and biotic consumption of oxygen and POC.

Biotic consumption was parameterised using literature-based ecophysiological rates combined with observed biomass distributions. Oxygen consumption rate (OCR in µmol $O_2$ g $AFDW^{-1}$ $h^{-1}$) and carbon assimilation expressed as POC in µg C assimilation µg $C^{-1}$ of biomass $m^{-1}$ $day^{-1}$ were compiled for hard substrate species. The observed species on the inside of monopile F1 were used as a representative group for the location. Where the video analysis identified higher order taxonomic groups, location-representative species for which relevant ecophysiological information was available were chosen (e.g. *Jassa herdmani* for Amphipoda).

Estimates of biomass were derived from the video footage. Conversion of OCR and carbon assimilation to AFDW was undertaken using conversion factors (Ricciardi & Bourget, 1998). Limited literature is available on biomass to organic carbon content conversion, especially for relevant individual species. Commonly used ranges in literature vary between 40-50 % (Kindeberg et al., 2024; Mackinson & Daskalov, 2007; Rueda et al., 2005; Wijsman et al., 1999). Therefore, the carbon content of the biomass (g AFDW $m^{-2}$ ) was determined using a conversion factor of 0.5 (Kindeberg et al., 2024) and subsequently applied to estimate the carbon assimilation rates of relevant species as input for the model.

A literature review produced information on the OCR for 10 out of 25 taxa found in monopile F1 and POC for four out of these 10 species (Harrington, 2010; Kumala et al., 2023; Mavraki, Degraer, Vanaverbeke, et al., 2020; Migné & Davoult, 1997; Noisette et al., 2016; Ortega et al., 1988; Pack et al., 2021; Souster et al., 2018; Voet et al., 2022). Seasonal or other variations were disregarded, as they were available for few of these species only. Subsequently, the OCR and POC values were used to arrive at a mean consumption and assimilation rate per mean biomass of carbon estimated per frame of the video analysis. Averages were derived from the available rates and assumed to represent the frame's species composition OCR and POC, in the absence of information of rates for every species present per frame.

The model reproduced the observed order of magnitude and general vertical structure of internal concentrations, including elevated values near the WRHs and reduced concentrations in zones above the upper and below the lower openings. While absolute agreement varied between monitoring days, the simulations consistently captured the dominant replenishment-controlled gradients observed inside the monopiles. Deviations between modelled and measured concentrations are attributed primarily to simplified and temporally constant boundary conditions for ambient oxygen and food availability, as well as uncertainties in biomass derived consumption rates. The evaluation indicates that the model is suitable for assessing relative patterns and sensitivities in internal water quality, rather than exact short-term concentration dynamics.
